# IgM^+^ and IgM^-^ memory B cells represent heterogeneous populations capable of producing class-switched antibodies and germinal center B cells upon re-challenge with *P. yoelii*

**DOI:** 10.1101/2021.03.04.433964

**Authors:** Susie L. Brown, Jonathan J. Bauer, Juhyung Lee, Enatha Ntirandekura, Jason S. Stumhofer

## Abstract

Memory B cells (MBCs) are essential for maintaining long-term humoral immunity to infectious organisms, including *Plasmodium*. MBCs are a heterogeneous population whose function can be dictated by isotype or expression of particular surface proteins. Here, aided by antigen-specific B-cell tetramers, MBC populations were evaluated to discern their phenotype and function in response to infection with a non-lethal strain of *P. yoelii*. Infection of mice with *P. yoelii* 17X resulted in the production of two predominant MBC populations: somatically hypermutated isotype-switched (IgM^-^) and IgM^+^ MBCs that co-expressed CD73 and CD80 that produced antigen-specific antibodies in response to secondary infection. Re-challenge experiments indicated that IgG-producing cells dominated the recall response over the induction of IgM-secreting cells, with both populations expanding with similar timing during the secondary response. Furthermore, using ZsGreen1 expression as a surrogate for activation-induced cytidine deaminase expression alongside CD73 and CD80 co-expression, ZsGreen1^+^CD73^+^CD80^+^IgM^+^ MBCs gave rise to class-switched IgG-producing plasmablasts that induced comparable titers of Ag-specific Abs as their IgM^-^ counterparts after adoptive transfer and infection with *P. yoelii*. Moreover, ZsGreen1^+^CD73^+^CD80^+^ IgM^+^ and IgM^-^ MBCs differentiated into B cells with a germinal center phenotype after adoptive transfer. A third population of B cells (ZsGreen1^-^CD73^-^CD80^-^IgM^-^) that emerges after infection responded poorly to reactivation in vitro and in vivo, indicating that these cells do not represent a population of MBCs. Together these data indicated that MBC function is not defined by immunoglobulin isotype, nor does co-expression of key surface markers limit the potential fate of MBCs after recall.

**Summary:** IgM^+^ and IgM^-^ MBCs that co-express CD73 and CD80 can differentiate into plasmablasts and GC B cells after re-challenge with *P. yoelii*.

## Introduction

Memory B cells (MBCs) represent a population of B cells that protect the host upon antigen (Ag) re-encounter. They can differentiate into antibody-secreting cells (ASCs) upon Ag recognition.^1^ is additional evidence that indicates that MBCs can re-enter the germinal center (GC) for further rounds of somatic hypermutation.^2–4^ While MBCs play an essential role in infection- and vaccine-induced protective immunity, many aspects of their generation, maintenance, and function in secondary responses remain vague. However, recent findings have shed new light on the biology of this cell type.^5^ Once thought of as a homogeneous cell population that expresses class-switched, somatically hypermutated B cell receptors (BCRs) generated within a GC, MBCs are instead a diverse heterogeneous population of cells with a GC-dependent and -independent origin.^6, 7^ Nevertheless, interpreting these recent findings on the biology of MBCs has proved to be complex. For instance, the ability of MBCs to re-enter the GC remains a point of controversy,^8–10^ as this event may be restricted to IgM^+^ MBCs and depend on the presence of persistent GCs.^3, 11^ Also, recent evidence indicates that a bottleneck event occurs that restricts the ability of MBCs to re-enter the GC.^12^ Conversely, others suggest that isotype-switched MBCs can re-enter GCs after challenge.^13^ In contrast, other evidence indicates that functionality is defined based on the presence or absence of cell surface markers, such as CD80, PD-L2, and CD73, rather than the isotype of the BCR.^2, 14, 15^ These findings suggest functional heterogeneity within the MBC pool. However, the contradicting results are likely the product of the models utilized to evaluate MBC function and highlight the need for additional studies to assess MBC heterogeneity on functional outcome, particularly in the context of infection-induced responses.

Studies in humans link *Plasmodium-*specific antibody (Ab) responses to protection from clinical disease.^16–20^ Additionally, experimental mouse models of malaria indicate that B cells and Abs are required for clearance of primary infections and protection against re-infection.^21–25^ Generation of long-lived plasma cells and MBCs during primary infection and their subsequent maintenance are needed for protection from secondary infections. However, the acquisition of Ab-mediated immunity against *Plasmodium* varies based on the number of exposures during each transmission season.^26, 27^ Although there is evidence in humans that MBCs and parasite-specific Abs can be maintained for long periods after infection,^28–33^ there is also evidence that a portion of MBCs generated against malarial antigens appear to be short-lived, and parasite-specific Ab titers drop rapidly with low re-infection rates.^34–38^ A similar drop in parasite-specific MBCs also occurs in mice over time after *P. chabaudi* infection.^39, 40^ These findings highlight many of the obstacles to developing an effective malaria vaccine.

MBCs are generated after *Plasmodium* infection in mice and humans; however, knowledge of their specificity, phenotype, origin, and affinity for malarial Ags is limited, although recent reports have provided novel insights into this subset of cells. For instance, MBCs produced after primary infection give rise to a faster secondary Ab response upon re-infection with *P. chabaudi*.^39–41^ This secondary response is initially dominated by the activation of IgM^+^ MBCs and is partially dependent on CD4^+^ T cells.^41^ Here, utilizing a *P. yoelii* infection model and B cell tetramers specific for two blood-stage Ags (MSP-1_42_ and AMA-1), we sought to evaluate the formation and diversity of the MBC compartment and determine which subpopulations of MBCs respond upon re-exposure to Ag. We found that MBCs that co-express CD73 and CD80 within the IgM^+^ and the isotype-switched (IgM^-^) MBC pools possess somatically hypermutated BCRs. In contrast, a population of IgM^-^CD73^-^CD80^-^ B cells that also emerge after infection display a BCR that contains few to no mutations, indicating a more germline state.

Furthermore, using a fluorescent ZsGreen1 protein as a surrogate for activation-induced cytidine deaminase (AID) expression and CD73^+^CD80^+^ co-expression, adoptive transfer of CD73^+^CD80^+^ZsGreen1^+^ IgM^+^ or IgM^-^ MBCs into naïve mice followed by infection with *P. yoelii* indicated that these MBC populations possess the ability to differentiate into plasmablasts or GC B cells. In contrast, CD73^-^CD80^-^ZsGreen1^-^ IgM^-^ B cells responded poorly to reactivation in vitro compared to their ZsGreen1^+^ counterparts, and they did not display the ability to differentiate into plasmablasts or GC B cells after adoptive transfer, indicating that they are not a functional MBC population. Collectively, these studies suggest that somatically hypermutated *P. yoelii*-specific CD73^+^CD80^+^ IgM^+^ and IgM^-^ MBCs respond to reactivation through cellular expansion and differentiation into IgG-producing plasmablasts and GC B cells.

## Materials and Methods

### Ethics statement

All animal studies described were done in concordance with the principles set forth by the Animal Welfare Act and the National Institutes of Health guidelines for the care and use of animals in biomedical research. All animal studies were reviewed and approved by the University of Arkansas for Medical Sciences Institutional Animal Care and Use Committee (Protocol number 3970).

### Mice

Male and female B6.Cg-Gt(ROSA)26Sor^tm 6 ( CAG- ZsGreen 1 ) Hze^/J (Ai6), B6.129P2- Aicda^tm( cre) Mnz^/J (*aicda*^cre^), B6.SJL-*Ptprc^a^Pepc^b^*/BoyJ (CD45.1) mice on a C57BL/6J background and WT C57BL/6J mice were obtained from The Jackson Laboratory. *Aicda^cre^*Ai6 reporter mice were generated by breeding homozygous *aicda^cre^* and Ai6 mice. Male BALB/c mice were obtained from Charles River Laboratories. All animals used in this study were between 7 and 12 weeks of age and were maintained and bred in specific pathogen-free facilities at the University of Arkansas for Medical Sciences in accordance with institutional guidelines.

### Parasites

Male BALB/c mice were infected with frozen parasite stock of murine *Plasmodium yoelii* 17X (MRA-749, BEI Resources Repository, Manassas, VA). Subsequently, 10^5^ parasitized erythrocytes derived from the passage were intraperitoneally (i.p.) injected into experimental groups of mice to establish infection. Mice were re-challenged at ≥ 90 days post-infection (p.i.) with 10^6^ parasitized erythrocytes, i.p. Male and female mice were used for the described experiments (except for passaging parasite stock). Primary and secondary infections were carried out in groups of mixed-sex mice. Adoptive transfer studies were performed separately for male and female mice with the transfer of cells from donor mice only into recipient mice of the same sex. During the primary infection, parasitemia was determined by flow cytometry as described,^42, 43^ or after re-challenge by Giemsa-stained thin-blood smears.

### Flow cytometry and antibodies

Single-cell suspensions were obtained by passing the spleen through a 40 μm cell strainer followed by lysis of erythrocytes by re-suspension in a 0.86% NH_4_Cl solution. Cells were maintained in complete RPMI (RPMI 1640 supplemented with 10% FBS, 10% sodium pyruvate, 10% non-essential amino acids, 10% L-glutamine, 10% Pen/Strep, and 1% β-mercaptoethanol). 3-5 × 10^6^ splenocytes from individual mice were aliquoted for staining into wells of a 96-well round-bottom plate. Cells were washed in FACS buffer (1 × PBS, 0.2% BSA and 0.2% 0.5M EDTA). Fc receptors were blocked in FACs buffer with anti-mouse CD16/32 (clone 24G2; BioXCell, Lebanon, NH) and normal mouse (ThermoFisher cat # 10410, Carlsbad, CA) and rat IgG (ThermoFisher cat # 10710C) in the presence of eF780 fixable viability dye (ThermoFisher cat # 65-0865-14). Surface staining was conducted with the following antibodies: CD73-PE-Cy7, GL- 7-AF488, PD-L2-PE, IgM-BV710, B220-AF700, CD80-BV605, CCR6-APC, CD38-AF700, CD19-BV650, CD19-BV510, B220-BV650, CD73-BV605, B220-BV785, CD138-BV711, CD80-BV650, streptavidin-BV510 (BioLegend), LIFR-PE (R&D Systems, Minneapolis, MN), CD38-eF450, CD39-PcP-Cy5.5, CD80-FITC, IgM-APC, GL-7-eF450, IgD-biotin, CD3e-APC- eF780, CD11b-APC-eF780, CD11c-APC-eF780, Ter119-APC-eF780 (ThermoFisher). For samples that did not require intracellular staining, cells were fixed with 4% paraformaldehyde. Fluorescence minus one (FMO) controls were used to set the positive gates and indicate background staining for histogram plots. Samples were acquired on an LSRIIFortessa (BD Biosciences, San Jose, CA) and analyzed using FlowJo version X software (BD Biosciences, Ashland, OR).

### Tetramer preparation

Recombinant His-tagged merozoite surface protein-1 (MSP-1_42_) (amino acids 1398-1754; Accession P13828) and apical merozoite antigen-1 (AMA-1) (amino acids 26-478; Accession AAC47193) proteins from *P. yoelii* 17X were produced in *E. coli* as previously described.^44, 45^ After purification and refolding the MSP-1_42_ and AMA-1 proteins, the His-tag was cleaved using a Thrombin kit (Millipore Sigma, St. Louis, MO). Confirmation of the His-tag removal was confirmed by W. blot using an anti-His Ab (ThermoFisher). MSP-1_42_ and AMA-1 were biotinylated and tetramerized with streptavidin-PE (SA-PE; Agilent, Santa Clara, CA) as previously described.^46^ A decoy reagent to gate out the non-MSP-1_42_ or AMA-1–specific B cells was constructed by conjugating SA-PE to DyLight 650 (ThermoFisher) followed by washing and removal of any unbound DyLight 650 and incubating with an excess of biotin as described.^46^

### Tetramer staining and enrichment

Single-cell suspensions of splenocytes were prepared as described above. Following red blood cell lysis, cells were resuspended in MACs buffer (Miltenyi BioTec, Bergisch Gladbach, Germany), and Fc receptors were blocked with anti-mouse CD16/32 (24G2; BioXCell) in a buffer containing normal mouse and rat IgG (ThermoFisher) for 20 minutes at 4°C. Then 10 nM of the prepared decoy reagent was incubated with the cells for 5 minutes at room temperature. MSP-1 or AMA-1–PE tetramers were then added at a concentration of 10 nM of SA, and the cells were incubated at 4°C for 30 minutes. Cells were subsequently washed in MACs buffer before incubating with anti-PE microbeads (Miltenyi) for 5 minutes at 4°C. Finally, cells were washed again in MACs buffer prior to positive selection on an AutoMACS Pro cell separator (Miltenyi). Recovered cells were surface stained and ran on a BD LSRIIFortessa. Collected data were analyzed using FlowJo version X software.

### BCR sequencing

MSP-1_42_ and AMA-1 tetramer-specific MBCs were sorted from day 90 *P. yoelii* or naïve *aicda*^cre^Ai6 reporter mice. Cells were pooled from the spleens of three individual mice then sub-gated based on the following parameters: live, single cells, decoy^-^tetramer^+^CD19^+^B220^+^ and sorted as follows: CD38^+^GL-7^-^IgM^+^ZsGreen1^-^ naïve B cells, CD38^+^GL-7^-^IgM^+^ZsGreen1^+^, CD38^+^GL-7^-^IgM^-^ZsGreen1^+^, and CD38^+^GL-7^-^IgM^-^ZsGreen1^-^ B cells. cDNA from individual B cells was used to generate a V(D)J library using a kit from 10x Genomics (Pleasanton, CA). Libraries were sequenced on a NextSeq 150-cycle high-output flow cell (Illumina, San Diego, CA). The Loupe V(D)J Browser (10x Genomics) was used to identify somatic mutations in the heavy and light chain genes of individual B cells. Individual V_H_ and V_L_ chain sequences were cross-checked and confirmed using the NCBI’s IgBLAST tool.

### Adoptive transfer experiments

*Aicda*^cre^Ai6 mice were sacrificed at least 70 days after infection with *P. yoelii* 17X parasites. Spleens were harvested and enriched for B cells by depleting other immune cell populations after labeling with biotinylated Abs against CD11b, CD3ε, Ter119, and CD11c followed by incubation with SA microbeads (Miltenyi Biotech) on an AutoMACS Pro cell separator (Miltenyi Biotech). Negative fractions were combined and stained to identify populations of MBCs. Live, single-cell CD19^+^B220^+^CD38^+^GL7^-^ MBCs were sorted based on the following parameters: IgM^+^ (CD73^+^CD80^+^ZsGreen1^+^IgG^-^IgD^+/-^), or IgM^-^ (CD73^+^CD80^+^ZsGreen1^+^IgM^-^IgD^-^ or CD73^-^CD80^-^ZsGreen1^-^IgM^-^IgD^+/-^) using a BD Biosciences FACSAria. FACS sorted cells were rested in cRPMI at 37°C for at least 30 minutes before prepping the cells for transfer. Cells were washed two times in sterile 1 × PBS and then resuspended in sterile 1 × PBS. Cells were loaded into a 1 ml syringe with a 30-gauge needle. Approximately 4-8 × 10^4^ CD73^+^CD80^+^ZsGreen1^+^IgM^+^, CD73^+^CD80^+^ZsGreen1^+^IgM^-^ or CD73^-^CD80^-^ZsGreen1^-^IgM^-^ B cells were transferred via the retro-orbital sinus into congenic CD45.1^+^ mice. Mice were challenged with 10^5^ *P. yoelii* pRBCs i.p. the following day. Transferred cells were recovered from infected mice on day 5, 8, or 28 p.i. by first staining with anti-CD45.2-PE and then enriched with the use of anti-PE microbeads (Miltenyi Biotech) followed by positive selection on an autoMACs Pro Separator.

### ELISAs

High binding Immunlon HBX plates (ThermoFisher) were coated with 2.5μg/ml of recombinant *P. yoelii* AMA-1 or MSP-1_42_ proteins in sodium bicarbonate buffer overnight at 4°C. Plates were blocked with 5% FBS (ThermoFisher) or FetalPlex (Gemini Bio, Calabasas, CA) in PBS for one hour at 37°C. Serum isolated from mice was initially diluted 1:50 or 1:00 and then serially diluted 1:3 down the plate leaving the last row for blocking buffer to serve as the blank control. HRP-conjugated IgG or IgM (Southern Biotech, Birmingham, AL) was incubated on the plate for 1 h at 37^°^C, and SureBlue substrate (Sera care, Milford, MA) was used for detection. The reactions were neutralized with stop solution (Sera care) before reading the plates. The plates were read on a FLUOStar Omega plate reader (BMG Labtech, Offenburg, Germany) at an absorbance of 450 nm.

### ELISpot assays

Following activation with 35% EtOH, ELISpot plates (Millipore cat # MAHAS4510) were coated overnight at 4°C with 2.5 μg/ml of *P. yoelii* recombinant MSP-1_42_ or AMA-1 protein in PBS. The following day, the plates were blocked for 30 minutes at room temperature with cRPMI. Afterward, 10^5^ splenocytes were plated and incubated overnight at 37°C in cRPMI. Next, plates were washed with PBS and then blocked with 5% FBS in PBS for 1 h at room temperature. Secondary IgM-AP (SouthernBiotech cat # 1020-04) or IgG-AP (SouthernBiotech cat # 1030-04) conjugated Abs were then incubated for 1 hour at 37°C in 5% FBS in PBS. Finally, an ELISpot developing reagent (consisting of BCIP and NBT; Sigma Millipore) was used to visualize spots; subsequently, an AID ELISpot reader system (Autoimmun Diagnostika, Strassberg, Germany) was utilized to calculate the number of ASCs.

For the MBC ELISpot assays, sorted populations of MBCs (CD73^+^CD80^+^ZsGreen1^+^IgG^-^ IgD^+/-^, CD73^-^CD80^-^ZsGreen1^-^IgG^-^IgD^+/-^, CD73^+^CD80^+^ZsGreen1^+^IgM^-^IgD^-^, or CD73^-^CD80^-^ZsGreen1^-^IgM^-^IgD^+/-^) were plated at a concentration of 5,000/well in 96-well round-bottom plates. MBCs were restimulated in the presence of the TLR7 agonist R848 (1 μg/ml; Invivogen, San Diego, CA) and human IL-2 (10 ng/ml; Peprotech, Rocky Hill, NJ) with or without splenic feeder cells depleted of B cells (50,000/well). Splenic feeder cells incubated with R848 and human IL-2 and MBCs incubated without restimulation or added splenic feeder cells served as controls. Cells were incubated in vitro for three days at 37°C. After three days, the cells were transferred to ELISpot plates pre-coated with 2.5 μg/ml of *P. yoelii* recombinant MSP-1_42_ and AMA-1 or 10 μg/ml anti-mouse Kappa (Southern Biotech). Cells were incubated for 16-24 h at 37°C. Spots were visualized as described above.

### Statistics

GraphPad Prism 9 (San Diego, CA) was used for all statistical analyses. Specific tests of statistical significance are detailed in figure legends.

## Results

### MBCs segregate into heterogeneous populations based on phenotype after *P. yoelii* infection

To phenotypically define MBC subsets after infection, C57BL/6 wild-type (WT) mice were infected with a non-lethal strain (17X) of *P. yoelii*. Periodically over the course of the infection and after clearance (Supplemental Fig. 1A), spleens were harvested and stained for several phenotypic markers associated with MBCs,^14, 15, 47, 48^ including proteins that regulate T cell activity (CD80, PD-L2), enzymes involved in the generation of adenosine from ATP (CD73, CD39), the chemokine receptor CCR6, and the receptor for leukemia inhibitory factor (LIFR). While the specific role of these proteins on MBC function is largely unknown, a loss of these proteins through germline deletion or inhibition of ligand-receptor interactions can negatively impact the immune system.^49–56^

As several effector B cell populations emerge after infection, we used a gating strategy to distinguish between CD138^+^CD38^lo/-^ plasmablasts (PBs), CD38^-^GL-7^+^ germinal center (GC) B cells, CD38^+^GL-7^+^ multipotent precursors,^6^ and expanding CD38^+^GL-7^-^ MBCs in the spleen. The CD38^+^GL-7^+^ multipotent precursors represent a transitional stage in B cell differentiation that precedes the formation of GC B cells and MBCs and possesses the ability to differentiate into these populations.^6^ Within the CD138^-^CD38^+^GL-7^-^ population B cells were further subdivided into IgM^+^ and IgM^-^ groups (Supplemental Figure 1B). It is important to recognize that a large proportion of the CD38^+^GL-7^-^IgM^+^ B cells are naïve B cells, as they are indistinguishable from MBCs based on this phenotype. Subsequent analysis of the MBC associated markers CD73 and CD80 revealed the segregation of IgM^+^ and IgM^-^ MBCs into two predominant populations: CD73^+^CD80^+^ and CD73^-^CD80^-^ (Fig. 1A). This CD73 and CD80 expression pattern resembled that seen for 4-hydroxy-3-nitrophenylacetyl (NP)-specific MBCs after immunization with NP-chicken γ-globulin (NP-CGG)^15^ or infection with *P. chabaudi*.^41^ Shortly after infection, expansion in the IgM^+^CD73^+^CD80^+^ population occurred, followed by a decline. Then a second expansion that peaked at day 70 post-infection (p.i.) was observed. Similarly, IgM^-^ CD73^+^CD80^+^ MBC numbers peaked at day 70. While IgM^-^CD73^+^CD80^+^ MBC numbers declined after day 70, the IgM^+^CD73^+^CD80^+^ MBCs were steadily maintained through day 350 p.i. (Fig. 1B). However, the decline in IgM^-^CD73^+^CD80^+^ cell numbers was not as steep as that seen for the IgM^-^CD73^-^CD80^-^ cells, which saw a >30-fold reduction in numbers from day 70 to day 350. Although, the overall number of IgM^-^CD73^-^CD80^-^ MBCs was higher in the spleen almost a year after infection than the IgM^-^CD73^+^CD80^+^ MBCs (Fig. 1B). The IgM^+^CD73^-^ CD80^-^ population encompasses MBCs and naïve B cells; it is not surprising that their numbers remained steady past day 70 p.i.

**Figure 1.**
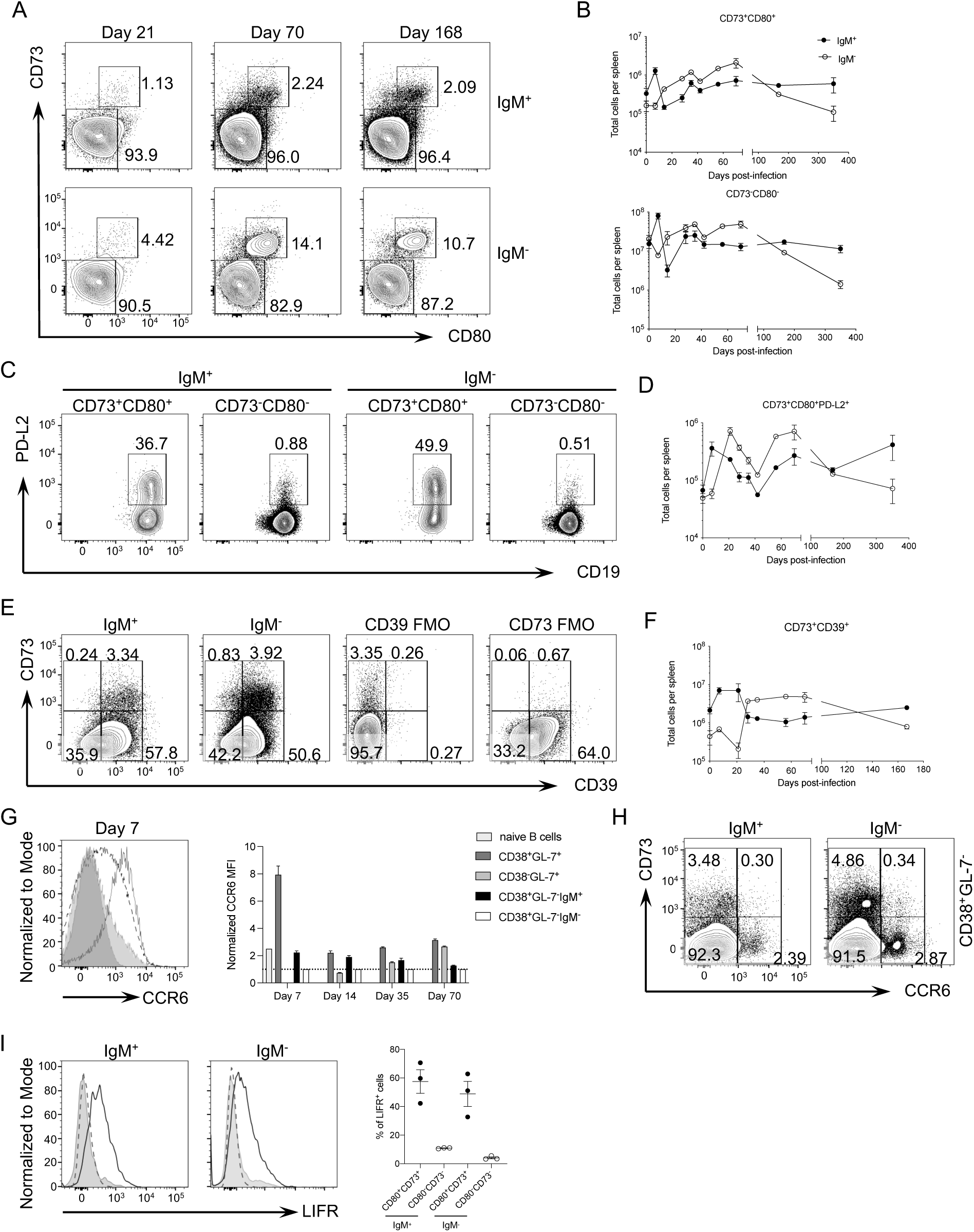
MBCs form a heterogeneous population after *P. yoelii* infection. (A-G) C57BL/6 mice were infected with *P. yoelii* 17X. To examine the expression of markers associated with MBCs, B cells were gated on live, single-cell CD138^-^CD19^+^B220^+^CD38^+^GL-7^-^ cells before looking at IgM expression. (A) Representative flow plots of CD73 and CD80 expression on IgM^+^ or IgM^-^ B cells at various times after *P. yoelii* infection. (B) Total number of IgM^+^ and IgM^-^ CD73^+^CD80^+^ and CD73^-^CD80^-^ B cells over 350 days p.i. (C) Representative flow plots of PD-L2 expression on IgM^+^ and IgM^-^ CD73^+^CD80^+^ and CD73^-^CD80^-^ B cells at day 167 p.i. (D) Total number of IgM^+^ and IgM^-^ CD73^+^CD80^+^PD-L2^+^ B cells over 350 days p.i. (E) Representative flow plots of CD39 and CD73 expression on IgM^+^ and IgM^-^ B cells at day 28 p.i. FMO = fluorescence minus one (F) Total number of IgM^+^ and IgM^-^ CD73^+^CD39^+^ B cells over 167 days p.i. (G) Representative histogram of CCR6 expression on CD38^+^GL-7^+^ B cells (solid black line), IgM^-^ (light gray shaded) and IgM^+^ (dotted black line) CD38^+^GL-7^-^ B cells at day 7 p.i. CCR6 FMO (dark gray shaded). Graph denoting CCR6 median fluorescence intensity (MFI) on splenic B cell populations at select times after *P. yoelii* infection. The CCR6 MFI for the CD38^+^GL-7^-^IgM^-^ B cells was assigned a value of 1.0 (dotted line), and the CCR6 MFI for all other populations was normalized to the CCR6 MFI of the CD38^+^GL-7^-^IgM^-^ B cells at each time point. (H) Representative flow plots of CCR6 expression on IgM^+^ and IgM^-^ CD38^+^GL-7^-^ B cells at day 28 p.i. (I) Representative histogram plots of LIFR expression on IgM^+^ and IgM^-^ CD73^+^CD80^+^ (solid black line) and CD73^-^CD80^-^ (dotted black line) B cells at day 28 p.i. LIFR FMO (gray shaded). Graph showing the frequency of LIFR expression on MBC populations at day 28 p.i. (B,D,F) Each data point shows the mean ± S.E.M. with 3-6 mice per time point. Representative of three independent experiments. (H) Error bars denote the mean ± S.E.M. Data are representative of three independent experiments with n = 3 mice per group. (I) Each point represents an individual mouse, and the error bars denote the mean ± S.E.M. Data are representative of three independent experiments with n = 3 mice per group.

Using the expression of CD80 and CD73 to define MBC populations, the expression of the other aforementioned markers associated with MBCs was examined. The majority of the CD73^-^ CD80^-^ cells, regardless of their isotype, did not express PD-L2. Also, PD-L2 expression was not uniform amongst the CD73^+^CD80^+^ cells; however, a higher frequency of IgM^-^CD73^+^CD80^+^ MBCs expressed PD-L2 compared to IgM^+^CD73^+^CD80^+^ MBCs (Fig. 1C). Regarding the expansion of the CD73^+^CD80^+^PD-L2^+^ MBCs, IgM^+^, and IgM^-^ populations displayed an early peak in expansion with the IgM^+^ cells peaking sooner (Fig. 1D). However, these peaks were followed by a contraction event in both populations through day 42 p.i., before each population expanded a second time. After this second peak plateaued at day 70 for the IgM^-^ cells, they contracted over time while the IgM^+^ cell numbers remained steady.

Since CD39 assists CD73 in breaking down ATP to adenosine,^57^ it was not surprising that most CD73 expressing MBCs (IgM^+^ and IgM^-^) co-expressed CD39. Also, a high frequency (>50%) of the CD73^-^ MBCs (IgM^+^ and IgM^-^) expressed CD39 (Fig. 1E). The IgM^+^ CD73^+^CD39^+^ cells peaked early after infection, then reduced between days 21 and 35 p.i. but thereafter, the numbers remained steady over time (Fig. 1F). In contrast, the IgM^-^ CD73^+^CD39^+^ cells plateaued at day 28 p.i. before declining in numbers after day 70.

The chemokine receptor CCR6 is associated with MBC precursors found within the light zone of the GC,^58^ and CD38^+^GL-7^+^ multipotent precursors express CCR6.^6^ Early on, after the infection, the per-cell expression of CCR6 was the highest for the CD38^+^GL-7^+^ multipotent precursors (Fig. 1G). As the infection progressed, the fluorescent intensity of CCR6 decreased on this population, correlating with the decline in this population (data not shown). Within the MBC pool, a small proportion of IgM^+^ and IgM^-^ MBCs showed expression of CCR6, with a higher expression on the IgM^+^ cells, which includes naïve B cells. Still, most CD73^+^ MBCs do not express this chemokine receptor, possibly suggesting that MBCs lose expression of this receptor with maturation or only a subset of them express it (Fig. 1G,H). Similarly, the expression of CCR6 was low amongst GC B cells upon their initial appearance but increased slightly over time (Fig. 1G). Signaling through the LIFR is linked to the self-renewal and survival of murine hematopoietic stem cells,^54, 59^ and it is upregulated and responsive to LIF stimulation on MBCs.^47^ Here, a sizable proportion of IgM^+^ and IgM^-^ CD73^+^CD80^+^ MBCs express the LIFR while very few CD73^-^CD80^-^ MBCs express this receptor after *P. yoelii* infection (Fig. 1I).

Expression of CD73 by B cells is thought to be associated with the expression of activation-induced cytidine deaminase (AID) through either a GC-dependent or -independent manner, as the majority of CD73^+^ MBCs have a high rate of somatic mutations in their BCR.^6, 14, 60^ To confirm that CD73 and CD80 co-expression coincides with AID expression, we crossed a mouse in which cre recombinase (*aicda*^cre/+^) replaces the endogenous AID protein with a mouse expressing the *R26*^ZsGreen1^ allele (Ai6). Thus, cre-mediated recombination in the subsequent offspring leads to permanent ZsGreen1 fluorescent protein marking upon expression of the *Aicda* locus (*aicda*^cre^Ai6 mice).^61, 62^ Regardless of isotype, the CD73^+^CD80^+^ MBCs favored expression of the ZsGreen1 protein, indicating they turned on AID transcription, whereas only a small fraction of IgM^+^ or IgM^-^ CD73^-^CD80^-^ MBCs expressed ZsGreen1 (Fig. 2). Thus, while CD73 and CD80 co-expression did largely correlate with AID expression, it was not absolute. Furthermore, while only a small proportion of IgM^+^ and IgM^-^ CD73^-^CD80^-^ B cells expressed ZsGreen1, sizable numbers of these cells were present in the spleen, particularly for those that expressed IgM, though the numbers for both populations were reduced on average compared to the CD73^+^CD80^+^ cells (Fig. 2C).

**Figure 2.**
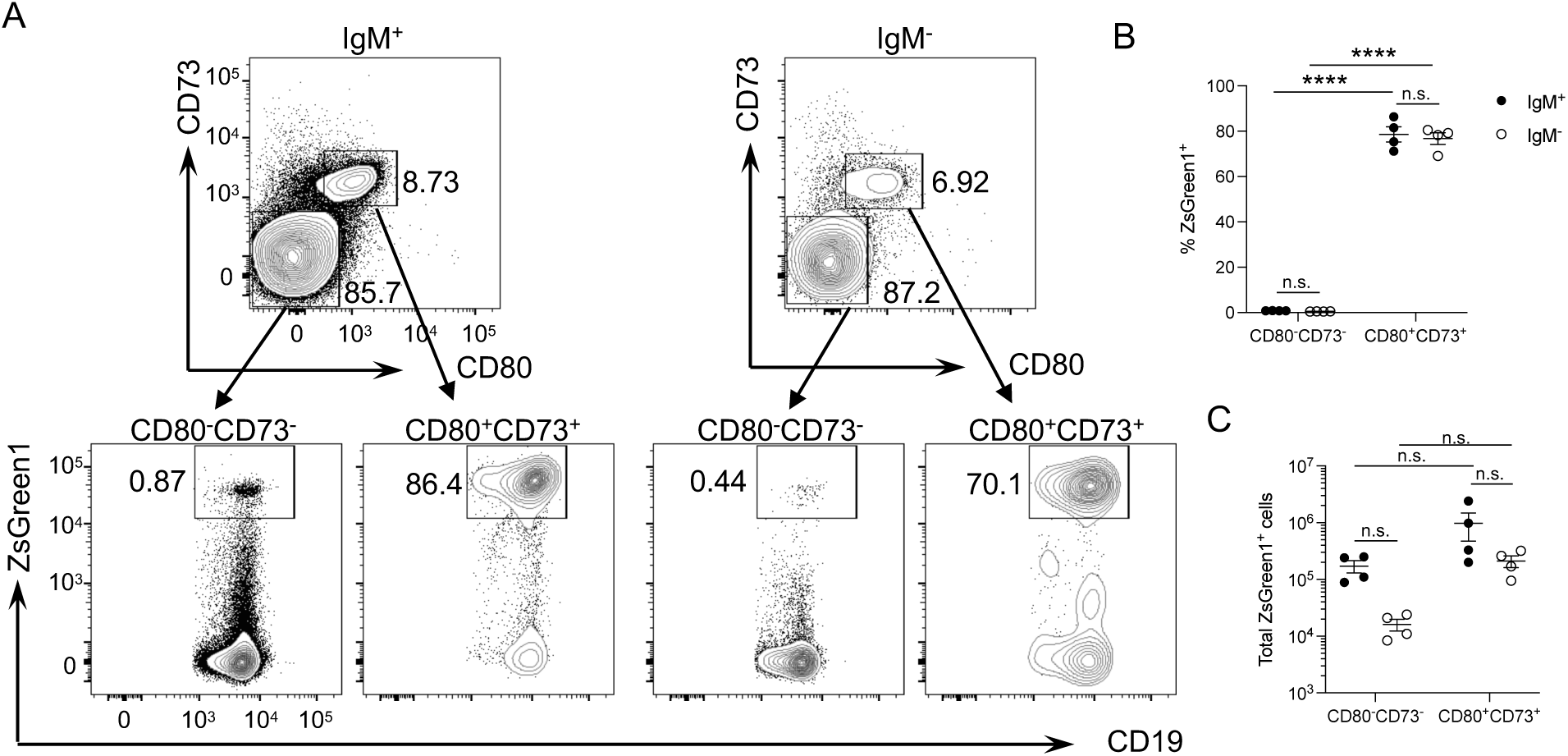
CD73^+^CD80^+^ MBCs predominantly express the ZsGreen1 protein, a surrogate marker for AID expression. *Aicda*^cre^Ai6 reporter mice were infected with *P. yoelii*. To examine ZsGreen1 expression, B cells were gated on live, single-cell CD138^-^CD19^+^B220^+^CD38^+^GL-7^-^ cells before looking at IgM expression. (A) Representative flow plots of ZsGreen1 expression in IgM^+^ and IgM^-^ CD73^-^CD80^-^ and CD73^+^CD80^+^ CD38^+^GL-7^-^ B cells at day 167 p.i. (B) Frequency and (C) total number of ZsGreen1^+^ IgM^+^ and IgM^-^ CD73^-^CD80^-^ and CD73^+^CD80^+^ CD38^+^GL-7^-^ B cells at day 167 p.i. Each point represents an individual mouse, and the error bars denote the mean ± S.E.M. Data are representative of three independent experiments with n = 3-4 mice per group. Significance was calculated by two-way ANOVA with post hoc Holm-Sidak’s multiple comparisons test. \**\**\*\* *p* < 0.0001, n.s. not significant.

Another layer of diversity in MBCs was recently described in the mouse model of *P. chabaudi* infection. A proportion of the IgM^+^ MBCs also expresses high amounts of IgD on their surface.^41^ Indeed, here, a sizable proportion of the IgM^+^CD73^+^CD80^+^ B cells were IgD^+^, whereas only a small proportion of the IgM^-^CD73^+^CD80^+^ were IgD^+^ (Supplemental Fig. 1C). Further phenotypic analysis indicated that most of the IgM^+^CD73^-^CD80^-^ B cells were positive for IgD expression, suggesting that these cells are predominantly naïve B cells. Similarly, the majority of the IgM^-^CD73^-^CD80^-^ B cells expressed IgD (Supplemental Fig. 1C). Since many IgM^-^CD73^-^CD80^-^ B cells do not express AID (based on the lack of ZsGreen1 fluorescence) and are IgM^lo/-^ and IgD^+^, these cells largely represent a population of B cells that have not undergone class switch recombination. Thus, it is unclear if this population represents an actual subset of MBCs. Lastly, we noted that the expression of the B-cell co-receptor component CD19 was significantly higher on IgM^+^ and IgM^-^ B cells that co-expressed CD73 and CD80 compared to those that did not express these cell surface markers (Supplemental Fig. 1C).

Together these data indicate that based on differences in the expression of several cell surface proteins, MBCs represent a population of cells, at least phenotypically, with multiple layers of diversity. However, the layers of diversity seem less dependent on the isotype of the BCR but rather dependent on the experiences of the B cell itself within the environment that forms after *P. yoelii* infection, as similar phenotypic populations are present in IgM^+^ and IgM^-^ MBC pools.

### Distinct MSP-1–specific plasmablasts, GC B cells, and MBCs are identifiable with a B-cell tetramer after infection

To substantiate our findings on MBCs from examining the total infection-induced B cell response, we utilized a B cell tetramer to examine Ag-specific B cells. Here, a phycoerythrin (PE)-conjugated B cell tetramer consisting of the 42 kD C-terminal end of the merozoite surface protein-1 (MSP-1) from *P. yoelii* was generated. This reagent was subsequently used to label Ag-specific B cells ex vivo, followed by magnetic bead-based enrichment of the labeled cells. As components of the tetramer itself, such as PE, can be bound by B cells, splenocytes were first stained with a decoy reagent to exclude them before incubation with the MSP-1 PE tetramer.^46^ Anti-PE-specific microbeads were then used to enrich the tetramer and decoy-specific B cells, which were subsequently labeled with Abs for analysis and quantitation by flow cytometry. After excluding dead cells, doublets, and non-B cells, B cell populations were identified and gated based on B220 and CD138 expression. Decoy^-^ MSP-1^+^ B cells were identifiable within this population before and after *P. yoelii* infection (Fig. 3A). Tracking MSP-1–specific B cells over one year after infection indicated that MSP-1–specific B cells rapidly expanded and peaked in cell numbers in the spleen shortly after infection and subsequently contracted by day 35. Thereafter a second but smaller expansion in cell numbers occurred and peaked by day 70 p.i. before contracting again by 100 days p.i. Hereafter the MSP-1–specific B cell numbers remained consistent over one year p.i., but the numbers were higher than those seen in naïve mice (Fig. 3B).

**Figure 3.**
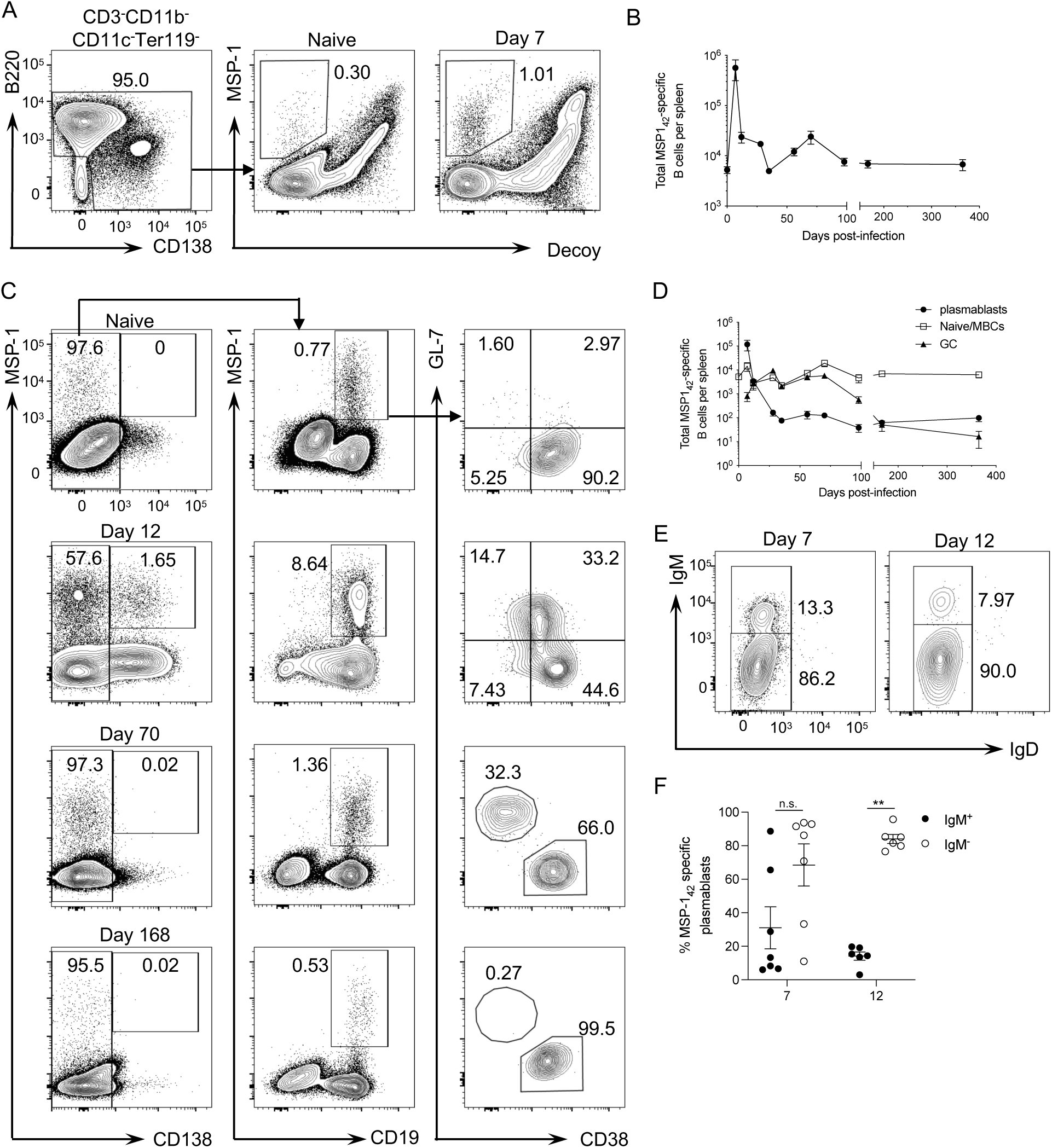
Identification and kinetics of MSP-1–specific B cells using a B cell tetramer. (A) Representative flow plot showing gated splenic B cell populations identified after excluding dead cells, doublets, non-B cells (CD3^+^CD11b^+^Ter119^+^), and enrichment with MSP-1 and decoy tetramers. Representative flow plots showing gating of MSP-1^+^decoy^-^ B cells from naïve and C57BL/6 mice infected for 7 days with *P. yoelii*. (B) Total number of MSP-1^+^ B cells over 365 days p.i. (C) Representative flow plots identifying CD138^+^MSP-1^+^ plasmablasts (left), CD138^-^ CD19^+^MSP-1^+^ B cells (middle), and CD38^+^GL-7^-^ (memory/naïve), CD38^+^GL-7^+^ (precursor), and CD38^-^GL-7^+^ (GC) B cells sub-gated from the CD138^-^CD19^+^MSP-1^+^ B cells (right). Splenic B cell populations were enriched with MSP-1 and decoy tetramers and identified after excluding dead cells, doublets, non-B cells (CD3^+^CD11b^+^Ter119^+^), and decoy^+^ cells. (D) Total number of MSP-1^+^ plasmablasts, memory/naïve, and GC B cells over 365 days p.i. (E) Representative flow plots displaying IgM and IgD expression by MSP-1^+^ plasmablasts at day 7 and 12 p.i. (F) The frequency of IgM^+^ and IgM^-^ MSP-1^+^ plasmablasts on days 7 and 12 p.i. in the spleen. (B,D,F) Each data point shows the mean ± S.E.M. with 6-15 mice per time point. (B,D) Data are combined from three independent experiments. (F) Data are representative of three independent experiments. A non-parametric Mann-Whitney t test calculated significance. \*\**p* < 0.01, n.s. not significant.

To corroborate our results with the MSP-1 tetramer, a similar set of experiments was performed using a B cell tetramer composed of another blood-stage Ag, apical merozoite antigen-1 (AMA-1) (Supplemental Fig. 2A,B). Rather than peaking at day 7 p.i. as the MSP-1^+^ B cells, AMA-1–specific B cells continued to expand in number throughout the infection, reaching their peak at day 28. Following a contraction phase, a second smaller wave of expansion occurred at day 70 before slowly contracting over time, a pattern that resembled that for the MSP-1^+^ B cells.

MSP-1–specific B cells possessed a naïve phenotype (CD38^+^GL-7^-^IgM^+^IgD^+^) in uninfected mice (Fig. 3C, *data not shown*), while distinct populations of effector B cells were noted in the spleen after infection, including plasmablasts, MBCs, and GC B cells (Fig. 3C). Also, MSP-1– specific CD38^+^GL-7^+^ precursor B cells^6^ were observed at days 7 and 12 after infection but were not maintained beyond this time (Fig.3C, *data not shown*). MSP-1–specific plasmablasts peaked in cell numbers by day 7 p.i. and contracted quickly by day 12 p.i. (Fig. 3D). By day 7, a transition from IgM^+^ to class-switched MSP-1–specific plasmablasts was evident (Fig. 3E,F), likewise for AMA-1–specific plasmablasts (Supplemental Fig. 2C,D). By day 12 p.i., the class-switched plasmablasts dominated the response (Fig. 3E,F). Whereas a distinct second wave of plasmablast formation started at 100 days after *P. chabaudi* infection and was visible out to 265 days p.i.,^41^ this degree of expansion was not apparent after *P. yoelii* infection (Fig. 3C, D). MSP-1–specific GC B cells peaked at day 28 p.i. and were still detectable at 100 days p.i., but by day 168, these cells were no longer prevalent. CD38^+^GL-7^-^ B cell numbers expanded at day 7 p.i. but contracted at day 12. Then this population went through another small expansion and contraction between day 28 and 35 p.i. before a large expansion took place after this time resulting in peak numbers by day 70 p.i. Afterward, this population contracted before leveling out by 24 weeks p.i. (Fig. 3C, D). Thus, the observed gains in cell numbers most likely represent MBC production waves, GC-independent at day 7 and GC-dependent through day 70 p.i., rather than an influx of naïve B cells.

### Tetramer-specific MBCs display a similar heterogenic phenotype as polyclonal MBCs

To determine if the MSP-1–specific MBCs phenotypically resembled the overall infection-induced MBC population, IgM^+^ and IgM^-^ tetramer^+^ MBCs were examined for expression of CD73 and CD80 throughout infection. Similar to findings in Figure 1, MSP-1–specific IgM^+^ and IgM^-^ MBCs were segregated into two populations: CD73^+^CD80^+^ and CD73^-^CD80^-^ over time (Fig. 4A). Few MSP-1–specific IgM^+^ and IgM^-^ cells upregulated CD73 by day 7 p.i., instead most of these CD38^+^GL-7^-^ B cells were predominantly CD73^-^CD80^-^ or CD80^+^ (Fig. 4B-C, *data not shown*). Following an initial expansion and contraction in cell numbers, MSP-1–specific IgM^+^ and IgM^-^ CD73^+^CD80^+^ and CD73^-^CD80^-^ MBCs peaked at day 70, after that contracting with time (Fig. 4B). Except for the MSP-1–specific IgM^-^ CD73^+^CD80^+^ MBCs, which were still declining, all other populations of MSP-1–specific MBCs had stabilized by one year p.i. (Fig.4B,C). This pattern of MSP-1–specific MBC expansion and contraction was in line with that observed for the overall MBC population after infection (Fig. 1).

**Figure 4.**
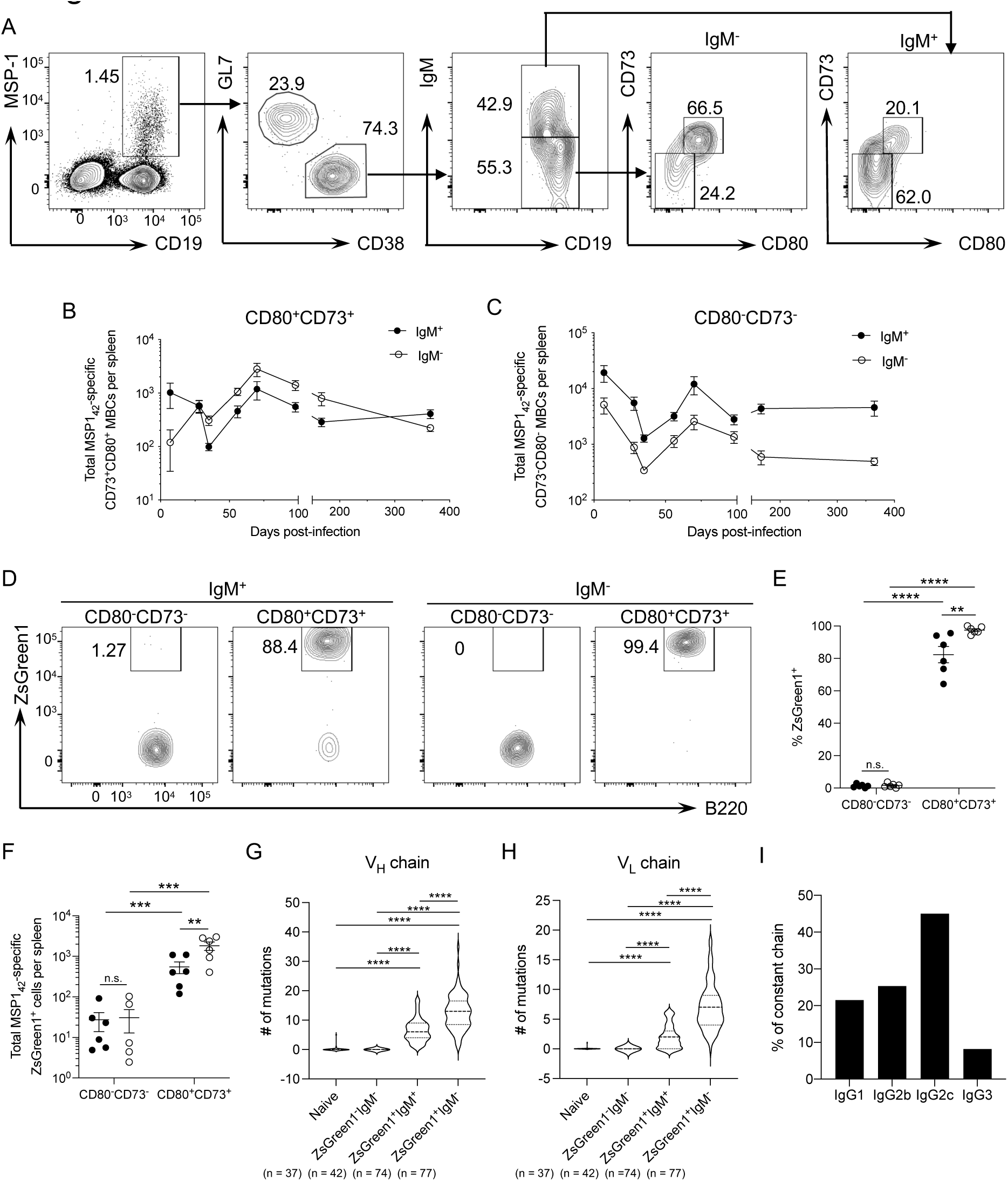
MSP-1–specific MBCs are phenotypically and genetically distinct populations. (A) Splenic B cell populations were enriched with MSP-1 and decoy tetramers and identified after excluding dead cells, doublets, non-B cells (CD3^+^CD11b^+^Ter119^+^), and decoy^+^ cells. Representative gating strategy used to identify MSP-1^+^ IgM^-^ and IgM^+^ MBCs based on CD73 and CD80 staining at day 70 p.i. Total number of MSP-1^+^ IgM^-^ or IgM^+^ (B) CD73^+^CD80^+^ or (C) CD73^-^CD80^-^ MBCs over 365 days p.i. Each data point shows the mean ± S.E.M. with 7-15 mice per time point. Data are combined from three independent experiments. (D) Representative flow plots showing the gates used to define ZsGreen1^+^ B cells within the MSP-1–specific IgM^+^ and IgM^-^ CD73^+^CD80^+^ and CD73^-^CD80^-^ MBC pools at day 100 p.i. (E) The frequency (F) and the total number of ZsGreen1^+^ B cells amongst the MSP-1–specific MBC populations at day 100 p.i. Each data point shows the mean ± S.E.M. with n = 6 mice. Data are representative of two independent experiments. Significance was calculated by two-way ANOVA with post hoc Holm-Sidak’s multiple comparisons test. \*\**p* < 0.01, \*\*\**p* < 0.001, \**\**\*\* *p* < 0.0001, n.s. not significant. Total number of mutations found in the CDR1 and CDR2 of the (G) heavy and (H) light chain gene of individual MSP-1–specific B cell clones. Numbers in parentheses represent the number of individual B cell clones assessed and graphed for each population. Dark dotted line in the violin plot represents the median, while the light dashed lines represent the quartiles. Significance was calculated by one-way ANOVA Kruskal-Wallis test with post hoc Dunn’s multiple comparisons test. **** *p* < 0.0001. (I) Frequency of IgG subclasses used by individually sequenced ZsGreen1^+^IgM^-^ MBC clones.

Examination of AMA-1–specific MBCs revealed some differences compared to the MSP-1–specific MBCs. While the pattern of expansion and contraction of the CD73^+^CD80^+^ MBCs looked similar, the frequency and number of AMA-1–specific IgM^+^ and IgM^-^ CD73^-^ CD80^-^ MBCs were lower than those observed for the MSP-1–specific cells, and their cell numbers were still declining a year after the infection had cleared (Supplemental Fig. 2E-G). Overall, examining the total infection-induced and Ag-specific MBC response after *P. yoelii* infection revealed that the spleen continues to generate IgM^+^ and IgM^-^ MBCs well after clearance of the infection. While CD73^+^CD80^+^ IgM^+^ MBC numbers reach a point of stability after infection, there is slow attrition in CD73^+^CD80^+^ IgM^-^ MBC numbers over time in the spleen.

Further examination of MSP-1–specific MBCs after *P. yoelii* infection indicated that most of the CD73^-^CD80^-^ MBCs expressed IgD, irrespective of IgM expression. Approximately half of the IgM^+^ CD73^+^CD80^+^ MBCs were IgD^+^. In contrast, the IgM^-^ CD73^+^CD80^+^ MBCs were primarily IgD^-^ (Supplemental Fig. 3). Examination of ZsGreen1 expression indicated that the MSP-1–specific IgM^-^ CD73^+^CD80^+^ MBCs were uniformly ZsGreen1^+^ with very few ZsGreen1^-^ cells present in this population, whereas there was more variability in ZsGreen1 expression in the IgM^+^ CD73^+^CD80^+^ population (Fig. 4D-F). Unlike the total infection-induced CD73^-^CD80^-^ B cells (Fig. 2C), little to no MSP-1–specific ZsGreen1^+^ cells accumulated in either the CD73^-^ CD80^-^ IgM^+^ or IgM^-^ pools. Lastly, the MBCs that co-expressed CD73 and CD80 also displayed a higher MFI for CD19 expression than the CD73^-^CD80^-^ B cells (Supplemental Fig. 3), a pattern that matched the overall infection-induced B cell response ( Supplemental Fig. 1). Hence, phenotypically distinct populations of Ag-specific MBCs exist in the spleen after infection with *P. yoelii*. However, they largely resemble the general infection-induced MBC cell populations and those generated in other mouse models of infection and immunization.

### *Plasmodium*-specific MBC subsets are genetically distinct

As B cells progress through the GC reaction, they accumulate mutations within their BCR, some of which increase their overall affinity for binding Ag. These high-affinity B cells can outcompete their brethren for T-cell help to maintain their survival; thus, allowing them to undergo further rounds of replication and somatic mutation or differentiate into long-lived MBCs and plasma cells. To explore the degree of somatic mutations in the BCR of various MBC populations after clearance of *P. yoelii* infection, individual heavy and light-chains of MSP-1– specific B cells (IgM^+^ and IgM^-^ ZsGreen1^+^, IgM^-^ ZsGreen1^-^) were sequenced from *aicda*^cre^Ai6 reporter mice. As shown in figure 4G-H, the IgM^-^ ZsGreen1^-^ B cells lacked mutations within their heavy and light chain genes. Furthermore, although these cells did not express IgM on their surface, they contained IgM transcripts. This finding supports the idea that these B cells are not the product of the GC and instead may represent naïve, immature, anergic, or recently activated B cells. They have downregulated IgM on their surface but have yet to undergo class switch recombination or somatic hypermutation, hence the lack of ZsGreen1 expression. The ZsGreen1^+^ IgM^-^ MSP-1–specific MBCs accumulated a significant number of somatic mutations within their V_H_ and V_L_ chains (Fig. 4G, H). Examination of their heavy chain constant region indicated that these MBCs favored expression of IgG2c, but a high proportion also expressed IgG1 or IgG2b (Fig. 4I). IgG3^+^ clones composed the smallest fraction of the ZsGreen1^+^ IgM^-^ clones sampled. While most of the ZsGreen1^+^IgM^+^ MBCs accumulated somatic mutations in their V_H_ or V_L_ chain genes, the number of mutations seen was significantly lower than the ZsGreen1^+^ IgM^-^ MSP-1–specific MBCs (Fig. 4G, H).

### MSP-1–specific MBCs rapidly expand after secondary infection

A hallmark of MBCs is their ability to differentiate into ASCs upon re-encounter with Ag. To determine how MBCs function during secondary infection, mice infected 90 days prior were re- challenged, i.p. with 10^6^ *P. yoelii* 17X pRBCs. MSP-1 tetramer^+^ B cells were examined on the day of re-challenge and 3, 5, and 9 days after re-infection (Fig. 5A). Expansion in the number of MSP-1^+^ B cells in the spleen occurred five days after re-challenge, and the numbers continued to rise over the next four days compared to the non-challenged mice (Fig. 5B). As expected, plasmablasts contributed to a large part of the early expansion in Ag-specific B cells after re- challenge. Their numbers expanded significantly over the first nine days after re-infection (Fig. 5A, C). An increase in the accumulation of MSP-1–specific IgM^+^ and IgM^-^ plasmablasts occurred at day 5 p.c., though more IgM^-^ plasmablasts were prevalent at this time (Fig. 5A, D). By day 9 p.c., the MSP-1–specific IgM^-^ plasmablasts dominated the response, though expansion was still occurring amongst the Ag-specific IgM^+^ plasmablasts. The MSP-1^+^ CD38^+^GL-7^-^ B cells, encompassing naïve and MBCs, and the MSP-1^+^ GC B cells also exhibited significant expansion by days 5 and 9 p.c. respectfully (Fig. 5C).

**Figure 5.**
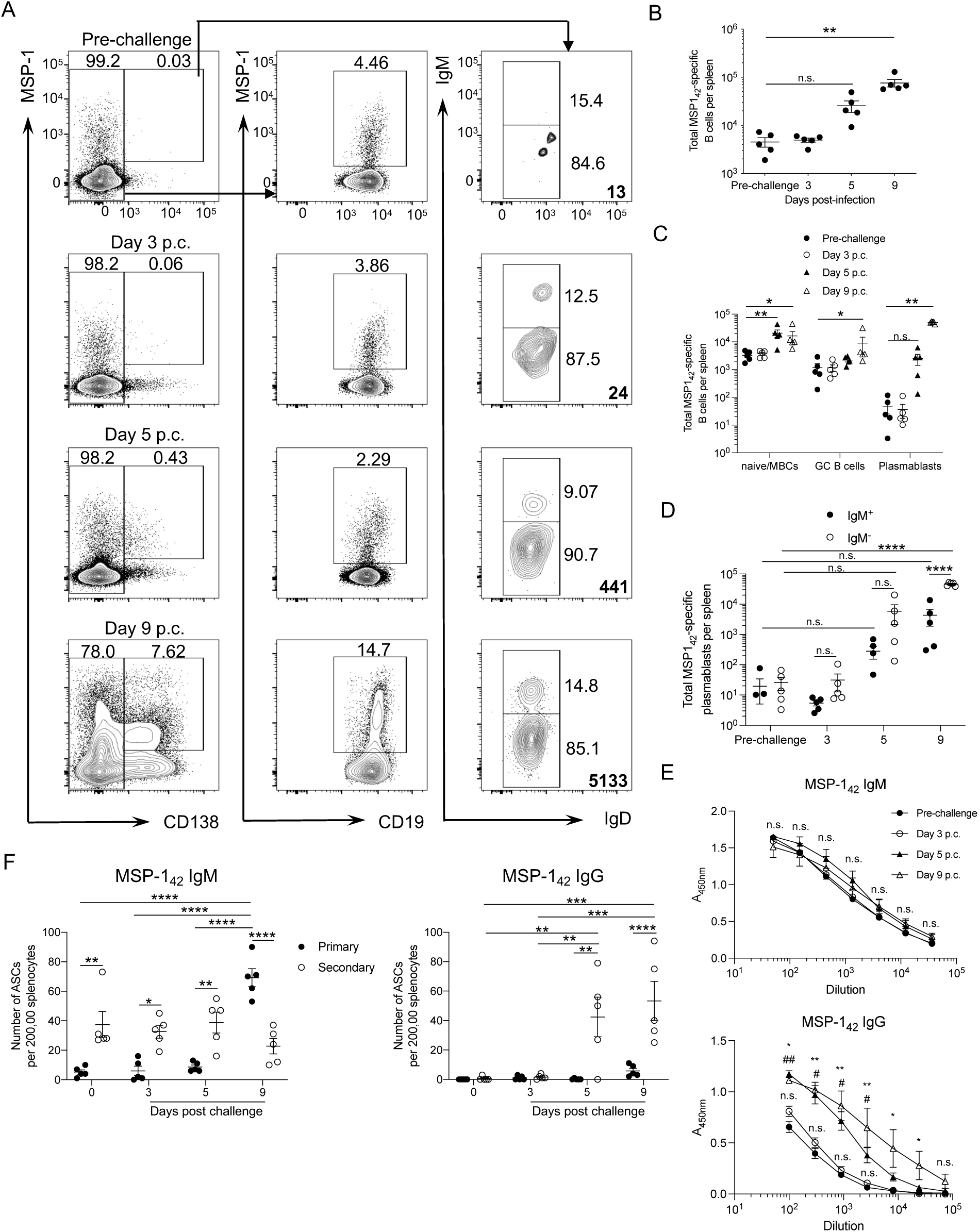
IgG^+^ ASCs dominate the secondary response to *P. yoelii*. Splenic B cell populations were enriched with MSP-1 and decoy tetramers and identified after excluding dead cells, doublets, non-B cells (CD3^+^CD11b^+^Ter119^+^), and decoy^+^ cells. (A) Representative flow plots identifying MSP-1–specific CD138^+^ plasmablasts, followed by sub-gating to look at IgM and IgD expression, and MSP-1–specific CD19^+^ B cells before and after re-challenge of mice at day 90 p.i. with 10^6^ *P. yoelii* infected RBCs. Bold numbers represent the number of cells on the plot, while non-bold numbers indicate frequency. (B) Total number of MSP-1^+^ B cells before and after re-challenge. C) Total number of MSP-1^+^ naïve/MBCs, GC B cells, and plasmablasts before and after re-challenge. (D) Total number of IgM^+^ and IgM^-^ MSP-1^+^-specific plasmablasts before and after re-challenge. (E) Serum IgM and IgG titers specific for MSP-1_42_ were determined by ELISA. (E) Splenocytes isolated before re-challenge (day 90) and post-challenge (days 3, 5, and 9) were used to determine the number of MSP-1-specific IgM and IgG ASCs by ELISpot. (B,C,D,F) Each point represents an individual mouse, and the error bars denote the mean ± S.E.M. with 5 mice per time point. Data are representative of three independent experiments. (B,C) Significance was calculated by one-way ANOVA Kruskal-Wallis test with post hoc Dunn’s multiple comparisons test. (D-F) Significance was calculated by two-way ANOVA with post hoc Holm-Sidak’s multiple comparisons test. \**p* < 0.05, \**\*p* < 0.01, \*\*\**p* < 0.001, \*\*\*\**p* < 0.0001, ^#^*p* < 0.05, ^##^*p* < 0.01, n.s. not significant. In **E,** the absorbance values for each dilution for the post-challenge samples were compared against the absorbance values for the pre-challenge samples. # represents the comparison between pre-challenge and day 5 p.c., and * represents the comparison between pre-challenge and day 9 p.c.

There was little to no change in MSP-1–specific IgM titers in the serum after re-challenge (Fig. 5E). In contrast, the MSP-1–specific IgG titers significantly increased by day 5 p.c., and they remained significantly elevated at day 9 (Fig. 5E). By ELISpot assay, there was only a minimal increase in MSP-1–specific IgM-secreting cells in some mice at day 5 p.c. (Fig. 5F), mirroring the flow cytometry results (Fig. 5D), but no further increase in numbers occurred at day 9 p.c. Alternatively, primary infected mice showed a significant expansion in MSP-1– specific IgM-secreting cells at day 9. In contrast, the MSP-1–specific IgG-secreting cells showed a significant expansion by day 5 p.c., and they remained elevated at day 9 (Fig. 5F), like the flow cytometry results (Fig. 5D). Furthermore, MSP-1–specific ASCs were not detectable in the spleen of mice with a primary *P. yoelii* infection until day 9 p.i. Their numbers were significantly lower than those observed in the re-challenged mice. Overall, these data indicate that the MBC response dominates the humoral response during secondary infection with *P. yoelii*. Together these results suggest that Ag-specific MBCs generated during primary infection with *P. yoelii* can differentiate into ASCs during secondary infection. Moreover, the IgG-producing ASCs dominate the secondary response to *P. yoelii* with only a marginal input by IgM-producing ASCs, which differs from the *P. chabaudi* re-challenge model findings.^41^ In comparison, we observed an expansion in B cells with a GC phenotype by day 9 p.c. However, it is unclear if MBCs are responsible for this increase.

### IgM^+^ and IgM^-^ *Plasmodium*-specific MBCs can differentiate into ASCs and GC B cells upon reactivation

The reduced expansion in Ag-specific IgM-secreting cells shortly after re-challenge indicates that the IgM^+^ MBCs may be non-responsive, possibly due to the high titers of MSP-1–specific Abs still present in the serum before re-challenge, or they have the capacity to class switch and become IgG-secreting cells after reactivation. A third possibility is that they preferentially proceed to the GC. To explore these possibilities, we used an adoptive transfer approach. Using *P. yoelii* infected *aicda*^cre^Ai6 reporter mice, IgM^+^ and IgM^-^ MBC populations expressing CD73, CD80, and the ZsGreen1 protein were sorted and adoptively transferred into naïve congenic mice that were subsequently infected with *P. yoelii* (Fig. 6A, Supplemental Fig. 4A,B). The functional capacity of the MBCs was evaluated 5, 8, and 28 days after infection.

**Figure 6.**
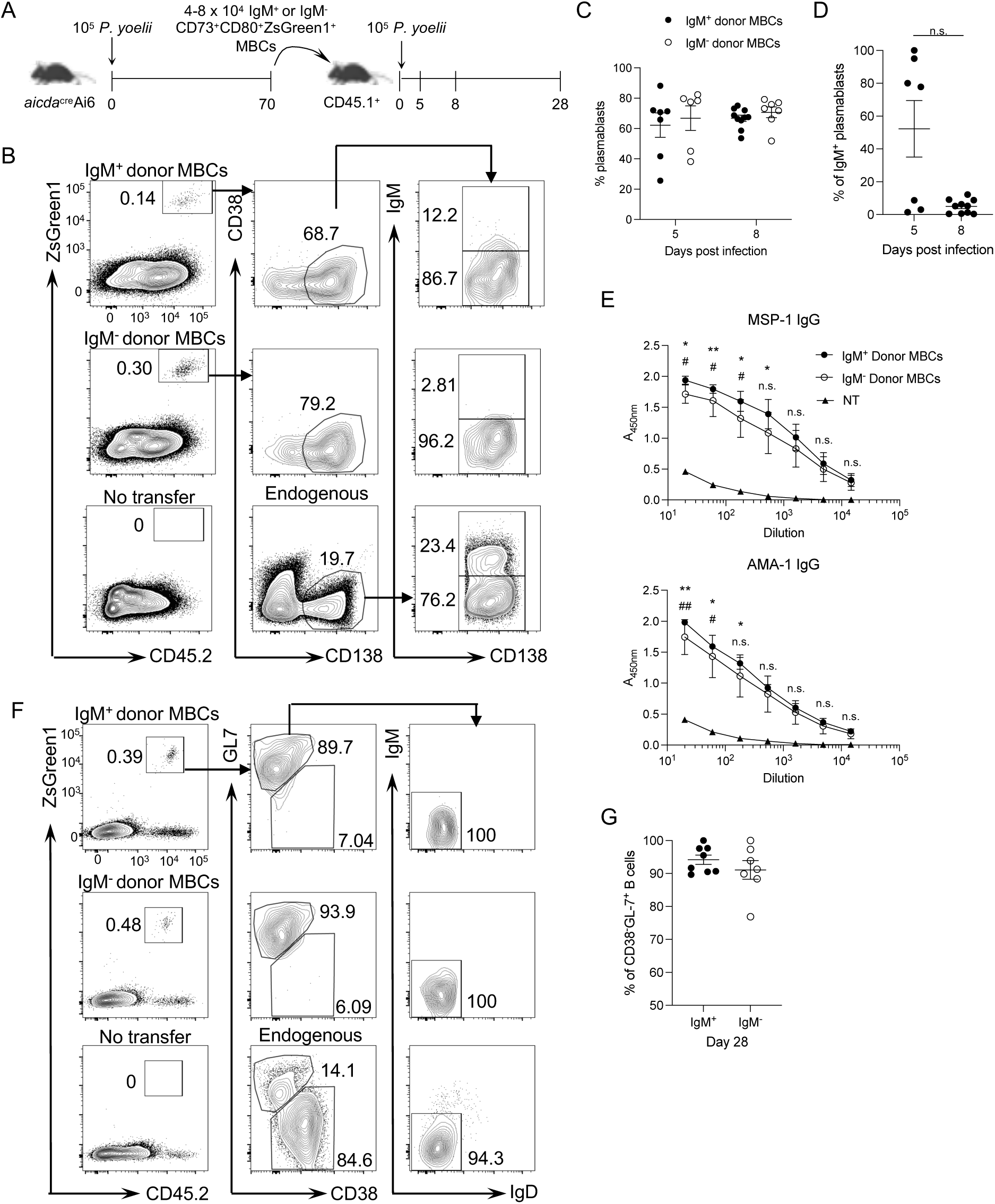
ZsGreen1^+^CD73^+^CD80^+^IgM^+^ MBCs predominantly differentiate into class- switched ASCs after reactivation. (A) Experimental model. CD73^+^CD80^+^ZsGreen1^+^ IgM^+^ or IgM^-^ MBCs were enriched and sorted from *aicda*^cre^Ai6 reporter mice ≥ 70 days after infection. 4-8 × 10^4^ MBCs were transferred into naïve congenic CD45.1^+^ mice that were challenged 24 h later with 10^5^ *P. yoelii* infected RBCs. Donor cells were recovered 5, 8, and 28 days after challenge. (B) Representative flow plots from day 8 p.i. showing gating strategy to identify CD45.2^+^ZsGreen1^+^ donor cells (left). Recovered donor cells were analyzed for their ability to differentiate into CD138^+^CD38^lo/-^ plasmablasts (middle) and the expression of IgM on the plasmablasts (right). Gates in middle and right plots based on the endogenous B cell population in the recipient mice. (C) Frequency of plasmablasts found within the recovered donor cell populations. (D) Frequency of IgM^+^ plasmablasts found within the recovered ZsGreen1^+^ plasmablast population. A non-parametric Mann Whitney t test calculated significance, n.s. not significant. (E) Serum IgG titers specific for MSP-1 and AMA-1 were determined by ELISA from day 8 infected mice. Samples were combined from three independent experiments. Significance was calculated by two-way ANOVA with post hoc Holm-Sidak’s multiple comparisons test. The absorbance values for each dilution for the transfer recipient samples were compared against the absorbance values for the no transfer recipient samples. * represents the comparison between IgM^-^ donors and no transfer and # represents the comparison between IgM^+^ donors and the no transfer. . \**p* < 0.05, \*\**p* < 0.01, #*p* < 0.05, ##*p* < 0.01, n.s. not significant (F) Representative flow plots from day 28 p.i. showing gating strategy to identify CD45.2^+^ZsGreen1^+^ donor cells (left). Recovered donor cells were analyzed for their ability to differentiate into CD38^-^GL-7^+^ GC B cells (middle) and the expression of IgM and IgD on the CD38^-^GL-7^+^ GC B cells (right). (G) Frequency of CD38^-^GL-7^+^ B cells found within the recovered ZsGreen1^+^ population derived from the donor IgM^+^ or IgM^-^ MBCs. (C,D) Each point represents an individual mouse, and the error bars denote the mean ± S.E.M. with 5-10 mice per time point. Data are pooled from two independent experiments. (G) Each point represents an individual mouse, and the error bars denote the mean ± S.E.M. with 6-8 mice per time point. Data are representative of three independent experiments.

The vast majority of the recovered donor cells at the early time points after infection displayed a CD38^lo/-^CD138^+^ phenotype indicative of plasmablasts (Fig. 6B,C), signifying that the transferred cells could differentiate into ASCs upon reactivation. Furthermore, while a proportion of the donor IgM^+^ MBCs differentiated into plasmablasts that expressed IgM, they began downregulating IgM expression as early as day 5 p.i., and by day 8 p.i. they were primarily IgM^-^ (Fig. 6B,D), suggesting their ability to undergo class switch recombination. This was the case as congenic mice that received IgM^+^ MBCs had detectable Ag-specific IgG in their serum by day 8 compared to congenic mice that did not receive a donor cell transfer (Fig. 6E). Thus, IgM^+^ and IgM^-^ MBCs primarily differentiate into ASCs, with the IgM^+^ MBCs undergoing class switch recombination upon reactivation.

Transferred donor cells were still identifiable at day 28 p.i. (Fig. 6F). These donor cells primarily displayed a GC B cell phenotype (CD38^-^GL-7^+^) at this time (Fig. 6F,G), indicating that IgM^+^ and IgM^-^ MBCs possess the ability to differentiate into GC B cells. Furthermore, Ag-specific MSP-1^+^ B cells were found amongst donor-derived GC B cells from each transferred MBC population (Supplemental Fig. 4C,D). Ag-specific IgG is readily detectable in the serum of congenic mice that did not receive donor MBCs at day 28 (Supplemental Fig. 4E). Therefore, the impact of the transferred IgM^+^ and IgM^-^ MBCs on Ab production was not distinguishable from the endogenous Ab response at this time. Overall, the transferred CD73^+^CD80^+^ZsGreen1^+^ IgM^+^ and IgM^-^ MBCs displayed a similar function upon recall, differentiating into ASCs early after reactivation and eventually progressing to a phenotype associated with GC B cell differentiation.

### IgM^-^ CD73^-^CD80^-^ MBCs are non-responsive to reactivation

While the above studies confirmed the function of the IgM^+^ and IgM^-^ CD73^+^CD80^+^ MBCs, they did not address the function of the IgM^-^ CD73^-^CD80^-^ B cells, the other potential MBC population identified in mice after *P. yoelii* infection (Fig. 1). To address the role of these B cells in a recall response, we attempted a similar approach, as shown in Figure 6. Instead, we sorted IgM^-^ B cells that were either CD73^+^CD80^+^ZsGreen1^+^ or CD73^-^CD80^-^ZsGreen1^-^ (Supplemental Fig. 4B) and adoptively transferred them into congenic mice that were challenged with *P. yoelii*. We hypothesized if these B cells are indeed MBCs, then like the IgM^-^ CD73^+^CD80^+^ZsGreen1^+^ MBCs, they should favor differentiation into plasmablasts upon reactivation. However, if they behaved identical to the CD80^-^PD-L2^-^ MBCs described in other systems,^2^ they should favor GC B cell development. In either scenario, they would potentially upregulate AID expression and hence express the ZsGreen1 fluorescent protein.

On day 8 p.i. cells recovered from congenic mice that received the IgM^-^ CD73^+^CD80^+^ZsGreen1^+^ donor B cells maintained ZsGreen1 expression. However, no ZsGreen1-expressing cells were recovered from the mice that received the IgM^-^ CD73^-^CD80^-^ ZsGreen1^-^ donor B cells (Fig. 7A), indicating that AID expression was not induced in the donor B cells. Furthermore, a negligible population of CD45.2^+^ cells that matched the background numbers recovered from the no transfer control group was isolated from the recipients of the IgM^-^ CD73^-^CD80^-^ZsGreen1^-^ donor B cells (Fig. 7B,C). Similar results were observed at day 28, as the number of recovered IgM^-^ CD73^-^CD80^-^ZsGreen1^-^ donor B cells matched the background amounts detected in the no transfer controls, and no ZsGreen1^+^ cells were recovered from these mice (Fig. 7D-F).

**Figure 7.**
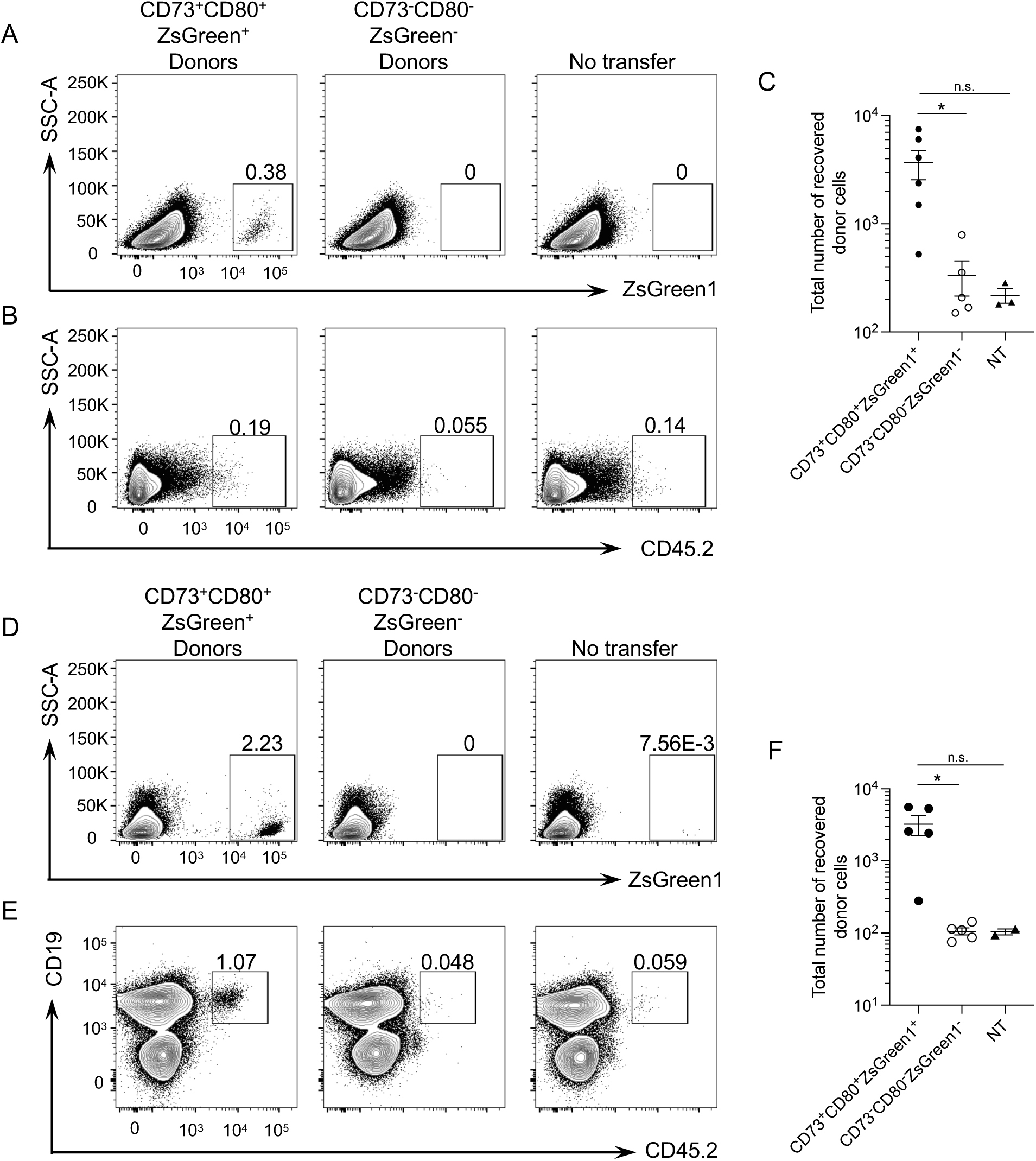
IgM^-^ CD73^-^CD80^-^ZsGreen1^-^ B cells fail to expand and upregulate ZsGreen1 expression after adoptive transfer and challenge with *P. yoelii*. (A) Representative flow plots showing the gating strategy to identify ZsGreen1^+^ cells derived from donor cells at day 8 p.i. (B) Representative flow plots showing the gating strategy to identify CD45.2^+^ cells derived from donor cells at day 8 p.i. (C) Total number of recovered donor cells at day 8 p.i. (D) Representative flow plots show the gating strategy to identify ZsGreen1^+^ cells derived from donors at day 28 p.i. (E) Representative flow plots showing the gating strategy to identify CD19^+^CD45.2^+^ B cells derived from donor cells at day 28 p.i. (F) Total number of recovered donor cells at day 28 p.i. (C, F) Each data point shows the mean ± S.E.M. with 2-6 mice per time point from two independent experiments. Significance was calculated by one-way ANOVA Kruskal-Wallis test with post hoc Dunn’s multiple comparisons test. \**p* < 0.05, n.s. not significant.

The significantly reduced yields seen for the cells derived from the IgM^-^ CD73^-^CD80^-^ ZsGreen1^-^ B cells may indicate that these cells have a reduced fitness upon transfer into a naive host, possibly due to competition with naïve B cells for a survival signal. Therefore, as a separate approach to evaluate their function, IgM^-^ CD73^-^CD80^-^ZsGreen1^-^ B cells, naïve, and IgM^+^ and IgM^-^ CD73^+^CD80^+^ZsGreen1^+^ B cells were sorted from *aicda*^cre^Ai6 reporter mice that recovered from a *P. yoelii* infection and restimulated in vitro. The IgM^+^ CD73^+^CD80^+^ZsGreen1^+^ B cells and naïve B cells could differentiate into IgM-producing ASCs, and some of the secreted IgM was Ag-specific (Fig. 8A,B). As seen in vivo, a proportion of the IgM^+^ CD73^+^CD80^+^ZsGreen1^+^ B cells could undergo class switch recombination after reactivation and differentiate into IgG- producing ASCs with some producing Ag-specific IgG (Fig. 8C,D). In contrast, the ability of naïve B cells to undergo class switch recombination and make IgG^+^ ASCs during this stimulation period was reduced compared to the IgM^+^ CD73^+^CD80^+^ZsGreen1^+^ MBCs. As expected, the ability of the IgM^-^ CD73^+^CD80^+^ZsGreen1^+^ MBCs to differentiate into IgG-producing ASCs was superior to the other B cell populations tested (Fig. 8C,D). The IgM^-^ CD73^-^CD80^-^ZsGreen1^-^ B cells responded poorly under the chosen in vitro stimulation conditions producing few to no IgG^+^ ASCs. Together this data indicates that IgM^-^ CD73^-^CD80^-^ZsGreen1^-^ B cells respond poorly to reactivation in vivo and in vitro, suggesting that these B cells are not a true population of MBCs.

**Figure 8.**
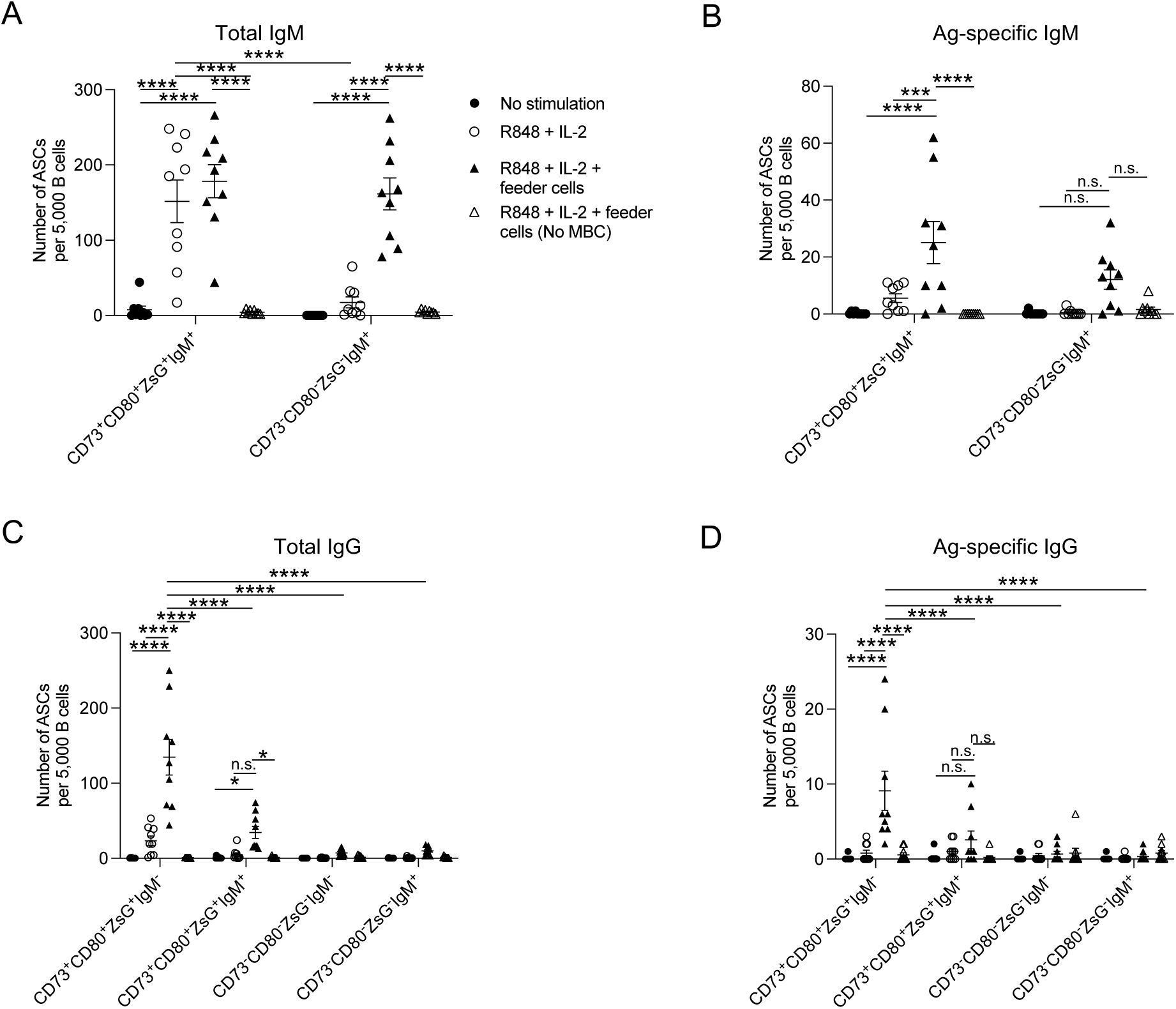
Stimulation of IgM^-^ CD80^-^CD73^-^ B cells in vitro does not induce their differentiation into ASCs. Splenocytes from *aicda*^cre^Ai6 reporter were isolated ≥ day 90 p.i to enrich and sort populations of MBCs based on the presence or absence of CD73, CD80, and ZsGreen1 expression and the presence or absence of IgM^+^ expression to determine their ability to differentiate into ASCs after restimulation in vitro. Sorted populations of MBCs were left unstimulated or restimulated with the TLR7 agonist R848 in the presence of recombinant IL-2 with or without B cell-depleted splenic feeder cells. (A) Total IgM^+^ or (B) IgM^+^ MSP-1/AMA- 1–specific ASCs and (C) total IgG^+^ or (D) IgG^+^ MSP-1/AMA-1–specific ASCs were determined by ELISpot. Each point represents an individual mouse, and the error bars denote the mean ± S.E.M. Data are representative of three combined experiments with n = 9 mice. Significance was calculated by two-way ANOVA with post hoc Holm-Sidak’s multiple comparisons test. \**p* < 0.05, \*\*\**p* < 0.001, \*\*\*\**p* < 0.0001, n.s. not significant.

## Discussion

Here, we indicate that MBC function upon recall with *P. yoelii* infection is not based on the isotype of the BCR, as IgM^+^ and IgM^-^ MBCs possessed the ability to differentiate into ASCs and GC B cells. These results contrast with the findings in other models that suggested that reactivation of IgM^-^ MBCs led predominantly to their differentiation into plasmablasts with very few producing GC B cells.^3, 4^ In both cases, mice were immunized with a non-replicating Ag. Hence, it is possible to speculate that the initial Ab burst due to the reactivation of transferred IgM^-^ MBCs resulted in the secretion of enough IgG to clear the Ag in these models, thus, preventing MBCs from differentiating into GC B cells. Support for this idea comes from the Pape et al. study, where the reactivation of IgM^-^ MBCs prevented the differentiation of endogenous B cells of the adoptive recipients from forming GCs.^4^ Here, we utilized a replicating parasite to investigate MBC function, which results in the maintenance of Ag well beyond clearance of the infection, as seen by the presence of Ag-specific GC B cells at least 70 days after resolution of the infection (Fig. 3D). Thus, our observation of IgM^-^ MBCs giving rise to B cells with a GC phenotype after transfer and challenge may partly be due to the inability of the Abs produced from MBC-derived plasmablasts to completely clear the parasite on their own. Thereby, a sufficient Ag load is available to allow for the accumulation of GC B cells.

Re-challenge of mice with *P. chabaudi* results in the rapid expansion of IgM-secreting plasmablasts, a proportion of which are T-cell independent, that peaks before the expansion of class-switched ASCs.^41^ While secondary infection with *P. yoelii* did induce an increase in IgM- secreting plasmablasts, it coincided with the class-switched ASC response, with the latter dominating the secondary response. The reduced expansion of IgM^+^ MBCs seen here with *P. yoelii* mirrors the results of secondary immunization with PE, which the authors attributed to the high serum quantities of anti-PE IgG circulating in these mice before re-exposure to PE.^4^ Similar to our findings here, the adoptive transfer of PE-specific IgM^+^ MBCs into naïve mice that were subsequently immunized with PE showed that these cells could recognize and respond to PE in an environment lacking circulating anti-PE Abs giving rise to IgM^-^ ASCs and GC B cells.^4^ Hence, we speculate that the ability of IgM^+^ MBCs to expand in response to secondary infection with *P. chabaudi* and not *P. yoelii* is due to differences in Ag-specific Ab quantity and/or quality in these mice at the time of re-infection. An idea supported by the differences in *P. yoelii* and *P. chabaudi* to establish secondary infections with blood parasitemia only detectable for the latter. *P. chabaudi* establishes a persistent infection after re-challenge, just as it did after primary infection but at a lower level of parasitemia.^63^

Thus, the Abs generated in response to primary infection with *P. yoelii* are more effective at preventing re-establishment of infection than those produced in response to *P. chabaudi* infection. Consequently, we speculate that critical epitopes recognized by the BCR of IgM^+^ MBCs and naïve B cells are preferentially bound by affinity-matured circulating Ag-specific Abs or high-affinity BCRs associated with IgM^-^ MBCs after re-infection with *P. yoelii*. In contrast, Ag is readily available for binding and reactivating IgM^+^ MBCs during secondary infection with *P. chabaudi*. The difference in the effectiveness of the Abs can be attributed to any number of factors, including increased titers and affinity of the Abs and differences in the production of Abs that target specific proteins essential for RBC binding and entry.

Importantly, our in vitro results indicate that the IgM^+^ MBCs can give rise to Ag-specific IgM-secreting plasmablasts, meaning that these MBCs can respond to restimulation. However, in vivo, they favored differentiation into IgM^-^ ASCs even without circulating Ag-specific Abs. Thus, additional factors such as inflammatory cytokines and additional signals derived from CD4^+^ T cells may influence differentiation and class-switching of IgM^+^ MBCs after reactivation in the absence of Ag-specific Abs.

Instead of favoring a model where MBC isotype expression dictates effector function, our results favor a model in which the expression of specific cell surface proteins defines their functional activity. In this case, co-expression of CD73^+^ and CD80^+^ expression by IgM^+^ and IgM^-^ MBCs represented a population of MBCs that had undergone extensive somatic hypermutation of their BCR and could differentiate into ASCs and GC B cells. The ability of the CD73^+^CD80^+^ MBCs to yield IgM^-^ ASCs during secondary responses echoed findings seen with CD80^+^PD-L2^+^ MBCs.^2^ However, these CD80^+^PD-L2^+^ MBCs did not give rise to GC B cells regardless of whether they expressed IgM or isotype-switched Ig, which differed from the results seen here. Instead, populations of IgM-expressing CD80^-^PD-L2^-^ or CD80^-^PD-L2^+^ MBCs preferentially gave rise to secondary GC B cells. While abundantly found after NP-CGG immunization,^2, 4^ MBCs with a CD80^-^PD-L2^+^ phenotype were rarely present in the spleen after *P. yoelii* infection regardless of isotype expression, as most PD-L2^+^ MBCs also expressed CD80 (Figure 1). Although IgM^+^ CD73^-^CD80^-^ MBCs were indistinguishable from naïve B cells in this model, the IgM^-^ CD73^-^CD80^-^ B cells found here did not respond to reactivation by differentiating into ASCs or GC B cells, indicating that these cells are not MBCs. The lack of somatic mutations in their BCR suggests that they are naive B cells that express IgD and low amounts of IgM. Another possibility is that these cells are anergic or in a transitional state following activation. Regardless of their origin or function, they are not a cell type contributing to the host’s immediate protection upon re-infection.

The cell markers used by immunologists to identify MBC populations suggest that phenotypical and functional heterogeneity exists within the MBC pool. However, their biological function on MBCs remains to be defined. For instance, the ectoenzymes CD39 and CD73 are surface proteins that convert ATP to AMP and adenosine. Whereas CD39 is expressed on naïve B cells,^49^ CD73 is upregulated on Ag-experienced and GC B cells,^6, 60^ and its expression is maintained on a subset of MBCs as seen here and by others.^2, 6, 14, 15, 41^ Furthermore, Tfh cells express high amounts of CD73,^50^ suggesting that the microenvironment of the GC is rich in ATP. However, its role in the GC is still unknown, as germline deletion of CD73 does not impair the GC response and MBC formation in response to NP-CGG immunization.^50^ Moreover, the mutational content of the BCR and MBC function in a recall response was not impacted by the loss of CD73. The only significant change observed in the absence of CD73 was reduced plasma cell accumulation in the bone marrow.^50^ Therefore, it is unclear why MBCs maintain expression of this enzyme on their cell surface. Although work with human naïve B cells and IgM^+^ MBCs showed that the ability of CD39 and CD73 to hydrolyze ATP to adenosine was required for class switch recombination.^64^ Furthermore, patients with common variable immunodeficiency with impaired class-switched Ab responses possess B cells that are selectively deficient in CD73 expression.^64^

Signaling through the LIFR receptor on hematopoietic stem cells is required for their survival;^54^ whether it promotes the survival of MBCs is unknown. However, the LIFR is functional on MBCs. They can respond to stimulation by LIFR agonists in vitro by inducing phosphorylation of Stat3.^47^ MBCs are long-lived and can self-renew; therefore, they possess attributes associated with hematopoietic stem cells. Another subset of B cells that can self-renewal is B1 B cells. Interestingly, a subset of CD25^+^ peritoneal B1 B cells expressed the LIFR and were responsive to stimulation by LIF.^65^ Thus, it is quite possible that LIF-LIFR signaling in MBCs plays a role in the maintenance of these cells, but further work is needed to address this possibility.

CCR6 expression is known to mark MBC precursors within the GC light zone; however, CCR6 expression is not required for MBC formation.^58^ Here, we found the highest expression of CCR6 on CD38^+^GL-7^+^ B cells early after infection with *P. yoelii*. Though a small proportion of GC B cells expressed CCR6, only a tiny subset of CD38^+^GL-7^-^ B cells also expressed this chemokine receptor, but they primarily did not co-express CD73. This result differs from the CD73^+^ MBCs found in the Peyer’s patch, where all IgA-bearing MBCs are positive for CCR6, and most express CD73 and PD-L2.^53^ CCR6 expression regulated the localization of MBCs within the Peyer’s Patch and promoted their survival. The lack of CCR6 expression on the CD73^+^ MBCs found here is intriguing given the additional finding that CCR6 expression is vital for the localization of MBCs in the spleen but not their generation or maintenance.^56^ Furthermore, altered localization of MBCs within the spleen impacts the ability of MBCs to be recalled to their cognate Ag in the absence of CCR6 expression, suggesting that the position of MBCs within the spleen is important for subsequent interactions with other cells such as T cells, which are implicated in the reactivation of some MBCs.^2, 66^ It is possible that infection with *P. yoelii,* as opposed to immunization with protein in adjuvant, leads to differences in CCR6 expression on MBCs observed in these models.

Although we found no evidence of rapid induction of IgM^+^ MBCs into IgM-secreting cells, this does not diminish the potential importance of these cells to the recall response in humans. In particular, Ag-specific IgM^+^ MBCs are identifiable in the blood of individuals from malaria-endemic regions,^38, 41, 67, 68^ and they can produce IgM Abs capable of neutralizing and promoting opsonization.^67, 68^ These Ag-specific IgM^+^ MBCs were more abundant than their IgM^-^ counterparts, even after years of repeated *Plasmodium* infections,^67^ suggesting their capacity to play a protective role from severe disease and illustrating the inefficient acquisition of protective IgG responses against *Plasmodium*. Although, the reduced rate of somatically mutated BCRs in IgM^+^ MBCs may make them better suited to recognize a broader range of variant *P. falciparum* Ags. Alternatively, the multimeric shape of IgM may improve its avidity for binding Ag and thus enhance its ability to bind and restrict parasite invasion of host cells.^68^

Lastly, as markers used to identify MBC populations continue to be defined and their function and biological role on MBCs is deduced, the challenge will be to determine if human MBCs similarly express these markers and if they display a similar functional role. Evidence of the expression of many of these markers on human MBCs is beginning to emerge. For instance, a recent study examining MBCs from patients with SAR-CoV-2 or acute *P. falciparum* malaria showed evidence of CD39 and CD73 expression by MBCs.^69^ However, the frequency of cells expressing CD39 and CD73 was reduced in all MBC populations in the malaria patients but not the COVID-19 patients. Given a described role for CD73 in lymphocyte homing,^70^ the homing mechanism of MBCs may be disrupted, or alternative receptors are used to promote homing into lymphoid tissues. It is also possible that given the link between CD73 and class-switching,^64, 71^ reduction in adenosine production affects the ability of these MBCs to class switch upon reactivation. Further understanding of the functional role of proteins expressed on MBC populations may provide valuable insight into why B cells have difficulty building protective memory against *P. falciparum*.

## Acknowledgments

*Plasmodium yoelii* 17X was obtained through BEI Resources, NIAID, NIH: MRA-749, contributed by D. Walliker. This work was supported by the Arkansas Biosciences Institute and National Institutes of Health (NIH) Grant AI116653 (J.S.S.) and P20-GM103625-Project 3 (to J.S.S.). In addition, the flow cytometry core is supported by the Translational Research Institute (Grant UL1-TR000039; NIH National Center for Research Resources and National Center for Advancing Translation Sciences) and the UAMS Center for Microbial Pathogenesis and Host Inflammatory Responses (Grant P20-GM103625; NIH National Institute of General Medical Sciences Centers of Biomedical Research Excellence). Special thanks to Andrea Harris for her technical assistance as part of the flow cytometry core.

## Author Contributions

Conceived and designed the experiments: JSS.

Performed the experiments: SLB, JJB, JL, EN.

Analyzed the data: SLB, JJB, JSS.

Wrote the paper: JSS.

---

**Supplemental Figure 1.**
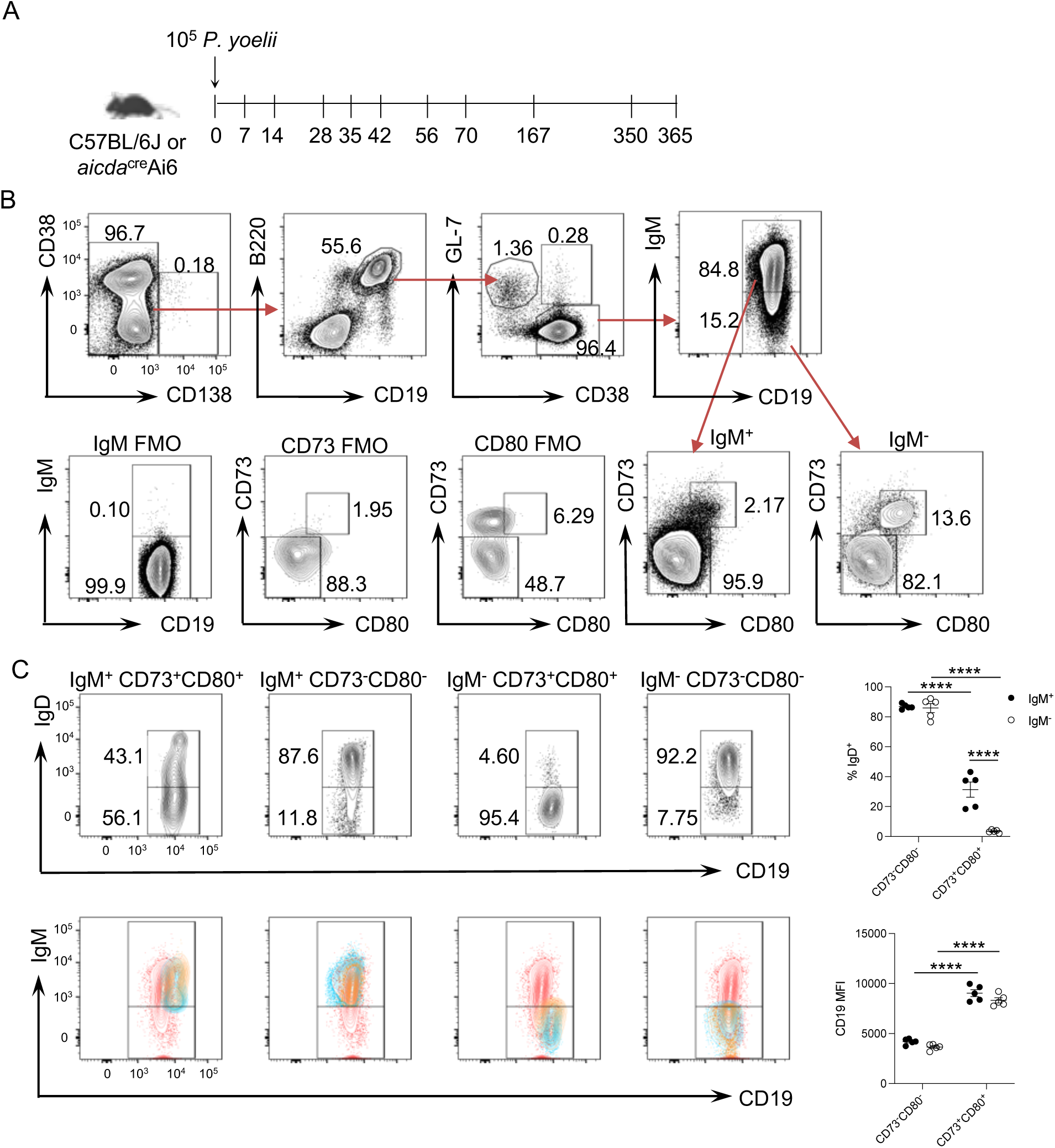
Gating strategy used to denote different B cell populations in the spleen. (A) Experimental model used in figures 1-4 and supplemental figures 1-3. C57BL/6 or *aicda*^cre^Ai6 reporter mice were infected with 10^5^ *P. yoelii* infected RBCs. Splenic B cell populations were analyzed by flow cytometry at indicated times p.i. after excluding dead cells, doublets, and non-B cells (CD3^+^CD11b^+^Ter119^+^). Experiments with tetramer enrichment also gated out decoy- cells before gating on B cell populations. (B) Representative flow plots from day 70 p.i. displaying the gating strategy used to identify IgM^+^ and IgM^-^ MBCs based on CD73 and CD80 expression patterns. Gates based on fluorescence minus one (FMO) controls. (C) Representative flow plots (upper) showing the gates used to distinguish IgD^+^ and IgD^lo/-^ B cells within the IgM^+^ and IgM^-^ CD73^+^CD80^+^ and CD73^-^CD80^-^ MBC pools at day 90 p.i. The graph (upper right) displays the frequency of IgD^+^ B cells amongst MBC populations at day 90 p.i. Representative flow plots (lower) displaying the IgD^+^ (orange) and IgD^lo/-^ (blue) MBCs from the gates in the upper plots overlayed over the total CD38^+^GL-7^-^ B cells (red) to illustrate the increase in CD19 expression by the CD73^+^CD80^+^ MBCs regardless of whether they express IgD. The graph (lower right) displays the MFI for CD19 expression for each population of MBCs. Each data point on the graphs represents an individual mouse, and the error bars denote the mean ± S.E.M. Data are representative of three independent experiments with 3-6 mice per time point. Significance calculated by two-way ANOVA with a post hoc Holm-Sidak’s multiple comparisons test. \*\*\*\**p* < 0.001.

**Supplemental Figure 2.**
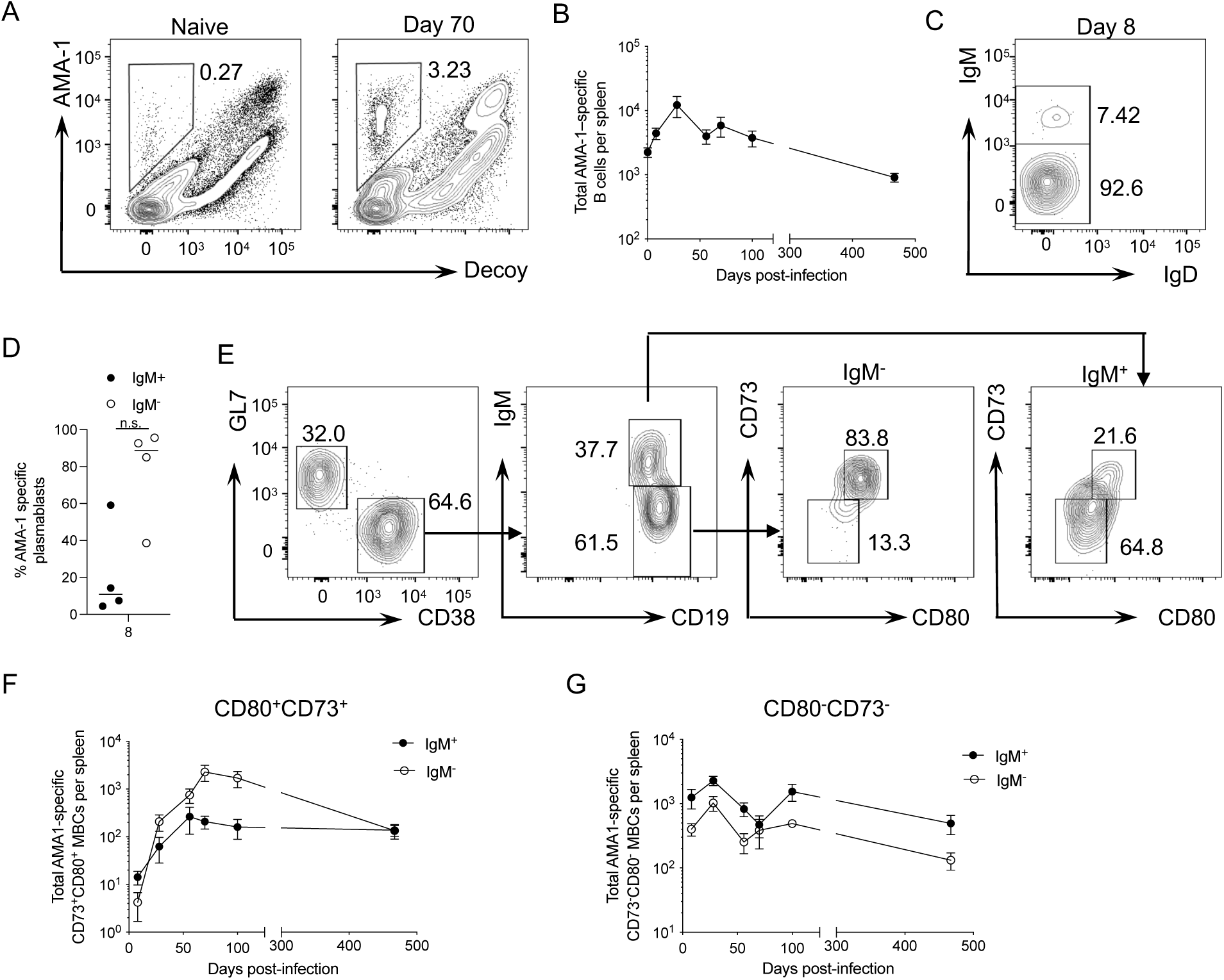
Detection and kinetics of AMA-1–specific B cells after *P. yoelii* infection. Representative flow plot showing gated AMA-1–specific B cells from naïve and day 70 infected C57BL/6 mice identified after excluding dead, doublets, and non-B cells (CD3^+^CD11b^+^Ter119^+^) and enrichment with MSP-1 and decoy tetramers. (B) The total number of AMA-1^+^ B cells over 467 days p.i. (C) Representative flow plots displaying IgM and IgD expression by AMA-1^+^ plasmablasts at day 8 p.i. (D) The frequency of IgM^+^ and IgM^-^ AMA-1^+^ plasmablasts at day 8 in the spleen. (E) Representative gating strategy used to identify AMA-1^+^ isotype switched and IgM^+^ MBCs based on CD73 and CD80 staining at day 56 p.i. Total number of AMA-1^+^ IgM^+^ or IgM^-^ (F) CD73^+^CD80^+^ or (G) CD73^-^CD80^-^ MBCs over 467 days p.i. Each data point shows the mean ± S.E.M. with 3-5 mice per time point from two independent experiments. Significance determined by a non-parametric Mann Whitney *t* test, n.s. not significant.

**Supplemental Figure 3.**
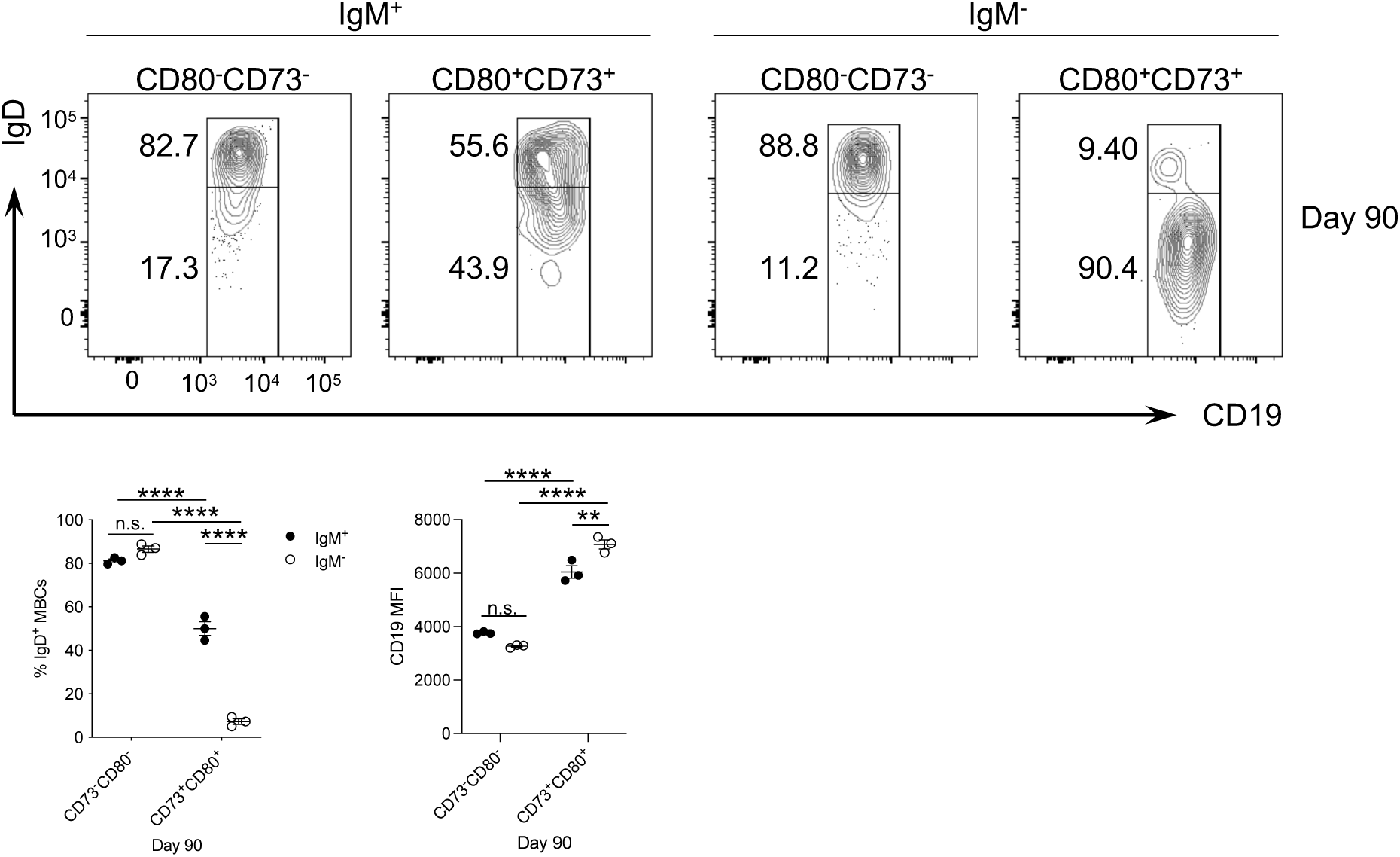
Expression of IgD on different MBC populations after *P. yoelii* infection. Representative flow plots showing the gates used to distinguish IgD^+^ and IgD^lo/-^ B cells within the MSP-1–specific IgM^+^ and IgM^-^ CD73^+^CD80^+^ and CD73^-^CD80^-^ MBC pools at day 90 p.i. The left graph displays the frequency of IgD^+^ MBCs amongst the MSP-1–specific MBC populations at day 90 p.i. The right graph displays the MFI for CD19 expression on the different MBC populations. Each data point represents an individual mouse and shows the mean ± S.E.M. with n = 3 mice. Data are representative of two independent experiments. Significance determined by two-way ANOVA with a post hoc Holm-Sidak’s multiple comparisons test. \*\**p* < 0.01, \*\*\*\**p* < 0.0001.

**Supplemental Figure 4.**
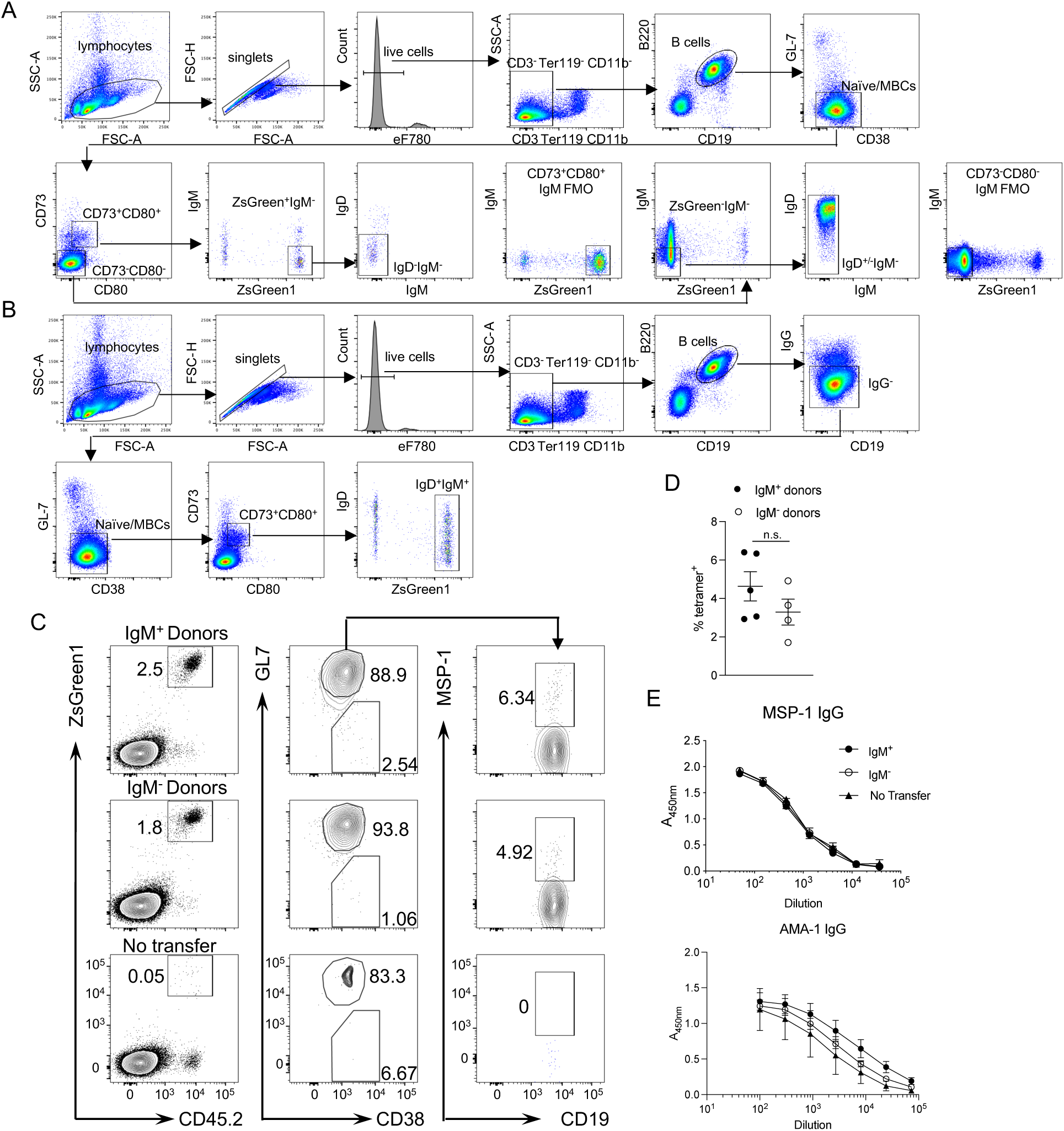
Gating strategy used to sort populations of MBCs from the spleen of *P. yoelii* infected mice. Strategy used to sort (A) IgM^-^IgD^-^ CD73^+^CD80^+^ZsGreen1^+^ or IgM^-^ IgD^+/-^ CD73^-^CD80^-^ZsGreen1^-^ B cells, or (B) IgM^+^IgD^+/-^ CD73^+^CD80^+^ZsGreen1^+^ B cells from *aicda*^cre^Ai6 reporter mice infected with 10^5^ *P. yoelii* infected RBCs for ≥ 70 days. Gates based on fluorescence minus one (FMO) controls. (C) Representative flow plots from day 28 p.i. showing the gating strategy to identify CD45.2^+^ZsGreen1^+^ donor cells (left). Recovered donor cells were analyzed for their ability to differentiate into CD38^-^GL-7^+^ GC B cells or retain their CD38^+^GL-7^-^ MBC phenotype (middle). Donor cells that displayed a GC phenotype were subsequently analyzed to determine their ability to bind MSP-1 (right). (D) Frequency of GC B cells that specifically bound MSP-1 within the recovered donor cells. Each data point on the graphs represents an individual mouse, and the error bars denote the mean ± S.E.M. Data are representative of two independent experiments with 3-5 mice. Significance calculated by two-way ANOVA with a post hoc Holm-Sidak’s multiple comparisons test, n.s., not significant. (E) ELISA from day 28 infected mice determined serum IgG titers specific for MSP-1 and AMA-1.

